# Bacteria control cell volume by coupling cell-surface expansion to dry-mass growth

**DOI:** 10.1101/769786

**Authors:** Enno R. Oldewurtel, Yuki Kitahara, Baptiste Cordier, Gizem Özbaykal, Sven van Teeffelen

## Abstract

Cells exhibit a high degree of intracellular crowding. To control the level of crowding during growth cells must increase their volumes in response to the accumulation of biomass. Using *Escherichia coli* as a model organism, we found that cells control cell volume indirectly, by increasing cell-surface area in proportion to biomass growth. Thus, dry-mass density, a readout of intracellular crowding, varies in proportion to the surface-to-volume ratio, both during the cell cycle and during perturbations such as nutrient shifts. On long time scales after shifts, initial dry-mass density is nearly restored by slow variations of the surface-to-mass ratio. Contrary to a long-standing paradigm, cell-envelope expansion is controlled independently of cell-wall synthesis but responds to the activity of cell-wall cleaving hydrolases. Finally, we observed rapid changes of Turgor pressure after nutrient shifts, which were likely responsible for initial changes of cell diameter and dry-mass-density. Together, our experiments reveal important regulatory relationships for cell volume and shape.

## Main text

A high level of macromolecular crowding of about 30-40% (*1, 2*) is essential for cellular physiology as it impacts processes ranging from macromolecular diffusion via protein-DNA interactions to protein translation (*3*). A robust readout for the level of crowding is the cellular dry-mass density, the ratio of dry mass (protein, RNA, etc.) to cell volume. Previous experiments in the model bacterium *Escherichia coli* suggest that dry-mass density is approximately constant (*4*–*6*). Therefore, cells are often assumed to increase their volumes in perfect proportionality to dry mass (*7, 8*). To increase volume, cells must expand their envelopes. Because the surface-to-volume ratio of rod-shaped bacteria changes in a width- and length-dependent manner (*S*/*V*= 4/[*W*(1 - *W*/3*L*)]), rates of surface and volume expansion differ during the cell cycle and during transitions between growth conditions. For density maintenance cells would thus need to tightly adjust their rate of surface growth or modulate width.

To study volume regulation, we developed a method to measure absolute single-cell volume *V*, dry mass *M*, and surface area *S* simultaneously and precisely during time-lapse microscopy. Volume and surface area are inferred from 2D cell contours obtained from phase-contrast or fluorescence images (Fig. 1A,S3). For dry-mass measurements we used Spatial Light Interference Microscopy (SLIM), a form of quantitative phase imaging (*9*) (Supplementary Note). The integrated optical phase shift of a cell is proportional to its dry mass (Fig 1B). To correct for cell-shape-dependent optical artifacts, we compared images to image simulations that are based on microscope parameters (*10, 11*).

**Figure 1.**
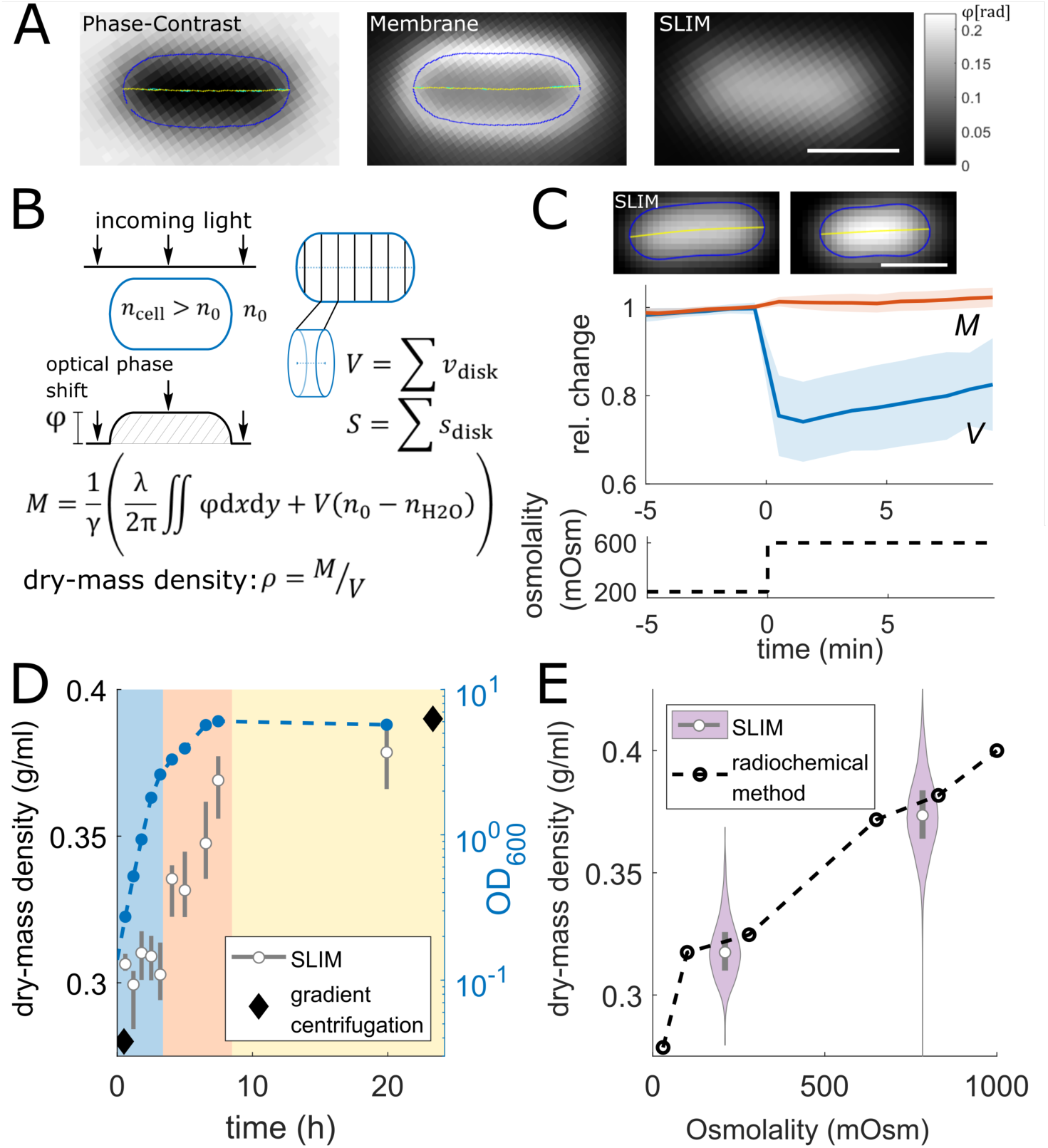
Measuring bacterial dry mass and volume using quantitative phase microscopy. **A:** Phase-contrast image (left), membrane stain (middle), and quantitative phase image (right) of WT cell (MG1655) grown in MM+mannose with cell contour (blue) and centerline (yellow). **B: Left:** Illustration of optical phase shift of light passing through a cell with refractive index *n*_cell_ > *n*_0_. **Right:** Volume and surface inference from calibrated 2D cell contour. Dry mass *M* equals the optical phase shift, corrected for *n*_0_, divided by the refraction increment *γ*. **C:** SLIM images before and after hyper-osmotic shock (top), relative changes of single-cell dry mass and volume (mean±std) during hyper-osmotic shock (middle), medium osmolality (bottom). **D:** Dry-mass density and optical density (OD600) during exponential growth (blue shade), during early (red) and late (yellow) stationary phase in LB. Comparison to values inferred from wet-mass densities (diamonds) (*2*). **F:** Dry-mass density increase with osmolality in MM+glucose, comparing SLIM and radiochemical free-water measurements (dashed line) (*3*). (grey rectangles=interquartile range; white circles=median; scale bars=1µm)

As a validation of our method, we found that dry mass remained constant after hyper-osmotic shock, demonstrating that mass measurements are independent of cell shape (Fig. 1C). Furthermore, mass density increased both with increasing medium osmolarity or upon entering stationary phase, in agreement with (*12*–*14*) (Fig. 1D,E). Mass density measurements were also confirmed through immersive refractometry (*15*) (Fig. S6).

We then applied our method to measure distributions of *M, V*, and dry-mass density *ρ* = *M*/*V* in two different *E. coli* strains (MG1655 and NCM3722) growing exponentially in different growth media of equal osmolarity (Fig. 2A,S7). Despite five-fold changes in average cell mass and volume (Fig. S7), the average density ⟨*ρ*⟩ varied by less than 15% between conditions – in a strain- and media-dependent manner (Fig. 2B). We confirmed these variations by refractive-index modulation (Fig. S8).

**Figure 2.**
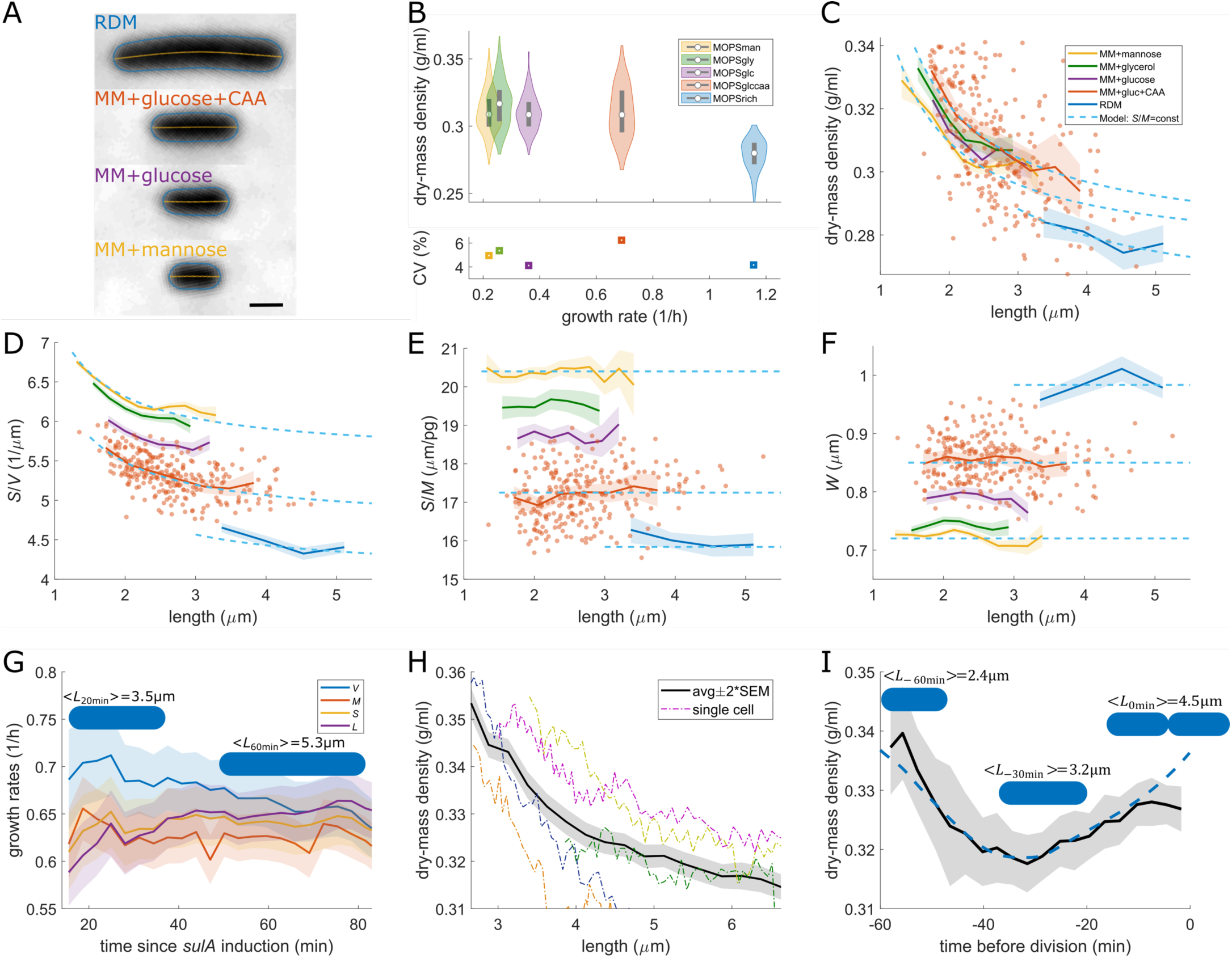
Cells increase cell surface but not cell volume in proportion to dry mass. **A-F:** Snapshots of non-dividing WT cells (MG1655) in different media at 30C. A: Phase-contrast images. B: Dry-mass density (white circles=median; grey rectangles=interquartile range) and coefficient of variation (CV). **C-F:** Length dependencies of dry-mass density (C), surface-to-volume ratio (D), surface-to-mass ratio (E), and width (F). **G-H:** Time lapse of filamenting cells (S290) in MM+glucose+CAA. G: Relative rates of volume, mass, surface, and length. H: Length dependency of dry-mass density. **I:** Time lapse of dividing cells (MG1655) in MM+glucose+CAA. Dry-mass density (solid line) and model (dashed line). (solid lines+shadings=average±2*S.E.M.; dots=single cell data)

Remarkably, cell-to-cell variations within conditions were smaller than 3-6% (CV) (Fig. 2B). Furthermore, up to half of the variations between non-dividing cells could be explained by a deterministic decrease of density as a function of cell length (Fig. 2C), which was also observed in time lapses of filamenting cells (Fig. 2H,S9) and through immersive refractometry (Fig. S10). To understand this seemingly complex dependency we investigated the possibility that cells might expand their surface areas rather than their volumes in direct proportion to biomass. To that end we decomposed mass density into surface-to-volume- and surface-to-mass ratios,

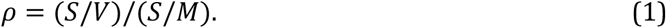

If cells grew in surface as they grow in mass, *S*/*M* should remain constant while density should vary in proportion to *S*/*V*. Indeed, *S*/*M* showed hardly any dependency on length (Fig. 2E). Furthermore, the length-dependent behavior of *ρ* was well described by the product of the average surface-to-mass ratio ⟨*S*/*M*⟩ and the surface-to-volume ratio of a model spherocylinder of width ⟨*W*⟩, where *W* is cell width. As a consequence, the volume growth rate *λ*_*V*_ = d(log *V*)/d*t* of short cells exceeded the constant mass-growth rate *λ*_*M*_ = d(log *M*)/d*t* by about 10% (Fig. 2G), in agreement with our prediction. In dividing cells, we then observed an increase of *ρ* towards the end of the cell cycle, due to the increase of ⟨*S*/*V*⟩ by septum formation (Fig. 2I).

While ⟨*S*/*M*⟩ is independent of length, it is strain- and media-dependent (Fig. 2E,S7). Specifically, the population-averaged ⟨*S*/*M*⟩ was nearly proportional to ⟨*S*/*V*⟩, such that variations of ⟨*ρ*⟩ between conditions remained small.

Next, we investigated how surface and mass were coupled during transitions between different growth conditions. First, we applied a nutrient upshift from a minimal medium containing mannose as carbon source (MM+mannose) to the same medium containing glucose and casamino acids (CAA) (MM+glucose+CAA) in batch culture and took regular single-cell snapshots (Fig 3A). Mass growth rate increased rapidly after the shift. Yet, ⟨*S*/*M*⟩ remained constant during about 50 min of growth, corresponding to 0.5 mass doublings. At the same time, mass density dropped by about 10% due to a similar decrease of *S*/*V*, largely due to changes of width. Only afterwards, ⟨*S*/*M*⟩ started to decrease, reaching its new steady-state value after ∼3.3 mass doublings. Therefore, cells change *S*/*V* and *S*/*M* on different time scales, which leads to transient and non-continuous changes of *ρ*.

**Figure 3.**
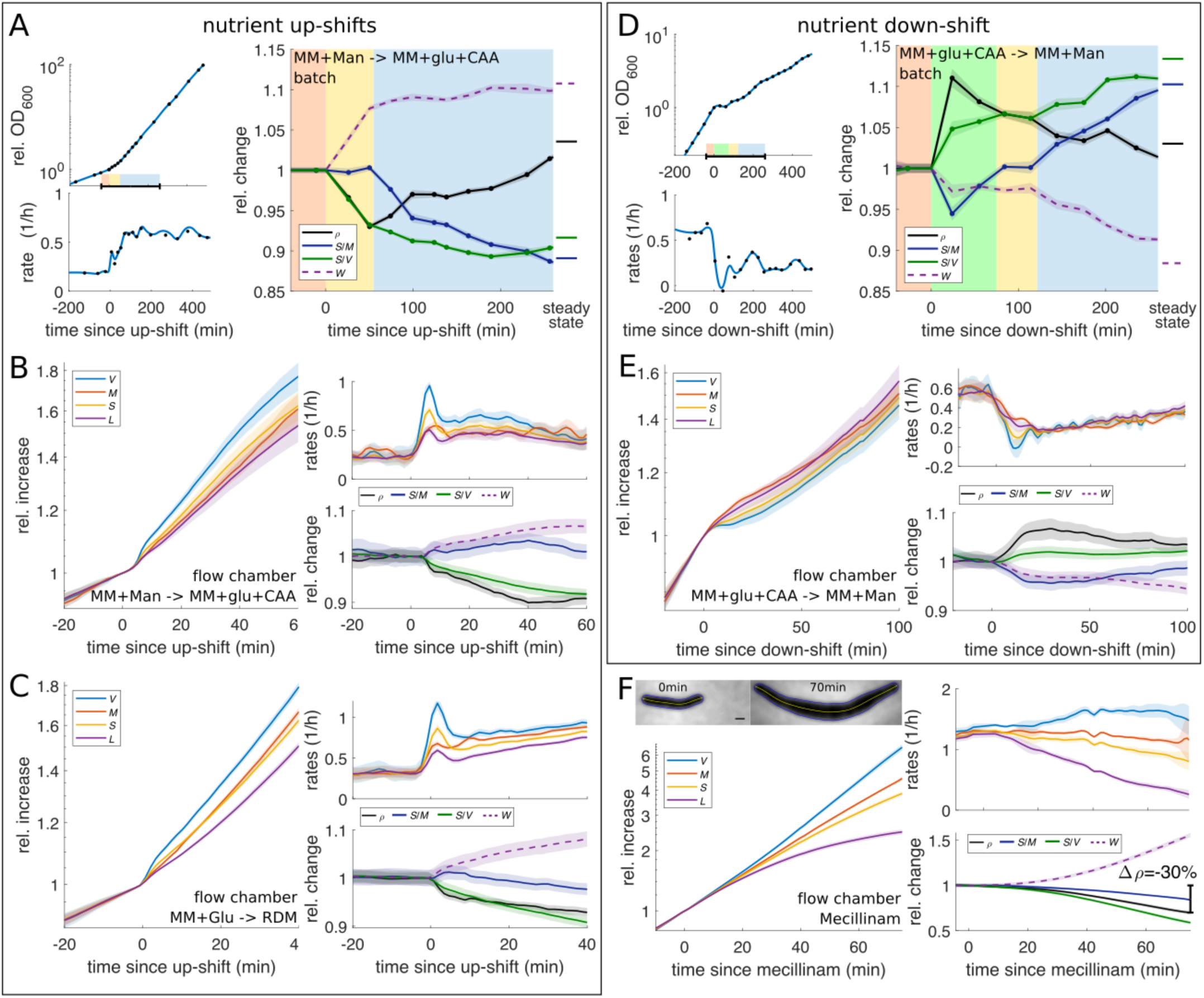
Shift experiments demonstrate robust surface-to-mass coupling despite variations of growth-rate, Turgor, and cell shape. **A:** Nutrient upshift from MM+mannose to MM+glucose+CAA in batch culture of WT cells (MG1655). Left: Growth curve and relative rate. Right: Relative changes of dry-mass density, surface-to-mass- and surface-to-volume ratios, and width from single-cell snapshots (right). Shaded background: *S*/*M* constant (yellow), *S*/*M* approaching new steady state (blue). **B:** Single-cell time lapse of the same nutrient shift as in A applied to filamenting cells (S290) in flow chamber. Relative increase (left) and single-cell rates (top-right) of volume, mass, surface, and length. Bottom-right: Same quantities as in A. **C:** Nutrient upshift from MM+glucose to RDM. Otherwise same conditions as in B. **D-E:** Nutrient downshift from MM+glucose+CAA to MM+mannose in batch culture (D) and flow chamber (E). Otherwise same conditions as in A, B. **F:** Mecillinam treatment in RDM. Otherwise same conditions as in B,C,E. Top-left: Snapshots. (solid lines+shadings=average±2*S.E.M.)

To follow single-cell behavior we applied the same shift to cells growing under the microscope, while inhibiting cell division (Fig. 3B,S13). During about 50 min after the shift, cell volume increased about 15% faster than cell mass, which led to a reduction of mass density by a similar amount as during the batch experiment. Despite slight variations of ⟨*S*/*M*⟩, possibly due to cell filamentation or surface attachment during microscopy, ⟨*S*/*M*⟩ was equal to its pre-shift value after 50 min, in agreement with the batch experiment. We observed the same behavior in a different shift from MM+glucose to RDM (Fig. 3C). Here, surface and mass grew at the same rate, while volume increased faster than mass and length increased more slowly, suggesting that not only volume but also length is indirectly controlled by surface and width. The dependency of length on surface and width is further supported by inverse single-cell correlations between rates of cell elongation and widening during steady-state growth (Fig. S12).

Interestingly, we observed striking deviations from the proportionality between mass and surface during the first few minutes after the shifts. Here, the rates of surface and volume expansion *λ*_*A*_ = d(log *A*)/d*t* and *λ*_*V*_ spiked due to rapid expansion of both cell length and width of about 1.5% each. This expansion is reminiscent of a hypo-osmotic shock (*16*), even though medium osmolarity was maintained constant. This observation suggests that Turgor pressures increased, and that initial surface expansion was largely elastic while the subsequent cell-diameter increase requires cell-wall remodeling. We observed a similar increase of cell dimensions when transiently increasing Turgor by continuously decreasing medium osmolarity (Fig. S14). In striking similarity to the nutrient upshift, these cells also showed a subsequent increase of cell diameter of about 10%, suggesting that cell-diameter changes during nutrient shifts might also be caused by Turgor.

We also investigated the dynamics of shape and mass during a corresponding nutrient downshift, both through time-lapse and batch-culture experiments (Fig. 3D,E). Similar to the upshift, we identified three phases. During the initial change of mass growth (<10 min), cells partially shrunk in volume due a sudden drop of cell diameter, demonstrating that osmotic pressure was reduced, even though medium osmolarity was maintained constant. We observed qualitatively similar but faster changes of Turgor upon depletion of glucose, using alpha-methylgucoside (alphaMG) (Fig. S15). After about 100 min (0.5 doublings) ⟨*S*/*M*⟩ had reached its pre-shift value (Fig. 3D,E), before increasing towards its new steady-state value. We therefore reasoned that the transient decrease of ⟨*S*/*M*⟩ was due to a concomitant reduction of Turgor, while surface material and dry mass might have increased at equal rates. Cell diameter and *S*/*M* reached their new steady-state-values after about 3.5h and 6h, respectively, consistent with the independence of *S*/*V* and *S*/*M* observed in the upshift experiment.

The robust coupling between surface and mass after sudden growth-rate changes gives support to a simple surface-growth law of the form

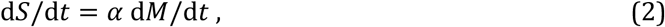

where alpha is a growth-condition-dependent coupling constant. During steady-state exponential growth *α*_ss_ = ⟨*S*/*M*⟩_ss_. Eq. (2) is mathematically equivalent to the equation d*S*/d*t* = *βM*, with *β* = *αλ*_*M*_. However, the former model is more appropriate, because *α* changes slowly during shifts, while *β* is nearly inversely proportional to *λ*_*M*_ (Fig. S16). Notably, our observations overturn the recent suggestion that surface grows in proportion to volume (d*S*/d*t* = *β*′*V*), and that cell diameter depends on *S*/*V* and a constant mass density (*7, 8*). Here, we found instead that *S*/*V* and *ρ* change according to independent variations of *W* and *S*/*M*.

Next, we tested whether surface-to-mass coupling was also robust with respect to non-physiological perturbations of cell dimensions. To that end, we inhibited the peptidoglycan (PG)-inserting multi-enzyme Rod complex responsible for rod-like cell shape, using the beta-lactam mecillinam (*17*) (Fig. 3F). Drug treatment led to rapid cell widening, as expected. Despite the severe perturbation, the rates of mass and surface growth remained nearly unaffected for about one mass mass-doubling time. As a consequence, dry-mass density dropped up to 30%, supporting our finding that cells grow in surface rather than volume.

Rod-complex inhibition is known to cause a strong reduction of the rate of PG insertion (*18, 19*). Therefore, our experiment demonstrates that surface expands independently of the rate of PG insertion, contrary to a major paradigm (*7, 20, 21*). To test whether surface expansion could proceed even upon arrest of cell-wall insertion, we treated cells with drugs that inhibit PG-precursor synthesis (D-cycloserine, Fosfomycin) or cross-linking (Vancomycin) (Fig. 4). For the latter, we used an outer-membrane-permeable *lptD*4213 mutant (*22, 23*). Cell-wall synthesis was severely reduced within 5-15 min, depending on the drug, according to the PG-synthesis-dependent rotation of MreB-actin (*24*) (Fig. 4A, Movies S1-3). Surprisingly, cells grew for an additional 15-40% in surface and mass at nearly unperturbed rates, up to the point of lysis (Fig. 4B). This finding was supported by an independent batch experiment (Fig. S17). Even when we combined Vancomycin treatment with a complex growth-rate shift, cell-envelope expansion proceeded normally (Fig. 4C). Therefore, PG synthesis is neither rate-limiting nor rate-determining for cell-wall expansion.

**Figure 4.**
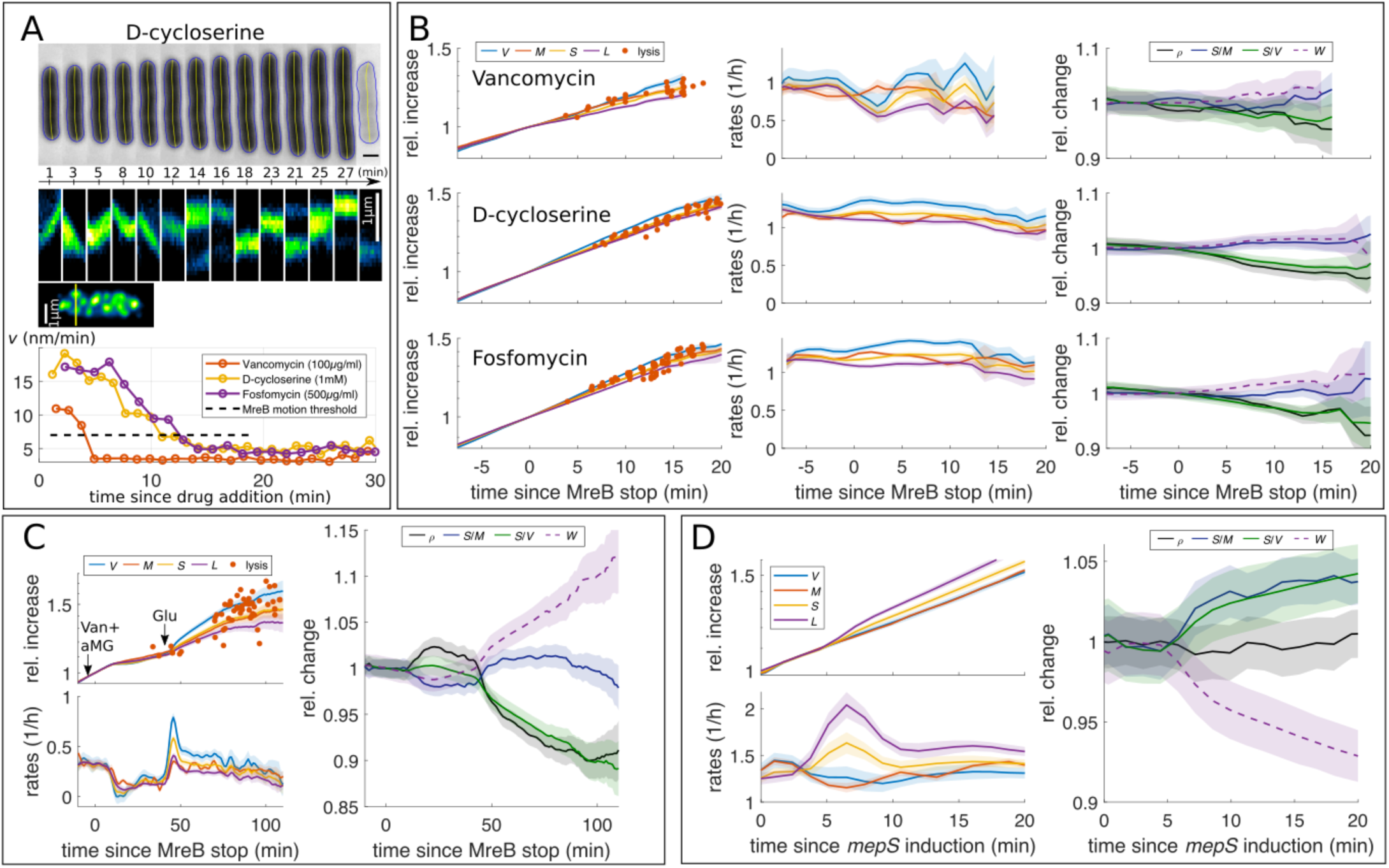
Surface expansion is independent of cell-wall synthesis. **A:** D-cycloserine leads to sudden lysis (top), only after MreB motion is arrested according to kymographs (middle) (S257). Average speed of MreB filaments after treatment with D-cycloserine, Fosfomycin (S257), and Vancomycin (S382). **B:** Growth after MreB-motion arrest is nearly unperturbed up to cell lysis. Red dots: single-cell mass at lysis. For description of lines see Fig 3B. **C:** Complex nutrient shift during Vancomycin treatment demonstrates surface-area control in the absence of cross-linking (S382). **D:** Overexpression of DD-endopeptidase MepS (B183) leads to transient increase of elongation and surface rates. (solid lines+shadings=average±2*S.E.M.)

Surface expansion during drug treatment requires the activity of lytic enzymes that cleave PG. Accordingly, we observed a rapid increase of the rate of surface expansion (Fig. 4D) upon overexpression of the DD-endopeptidase MepS, which cleaves cell-wall crosslinks (*25*). However, surface rate increased only transiently (∼5 min), demonstrating that enzyme activity is tightly regulated at the molecular level.

Together, we found a robust coupling between surface area increase and dry mass growth. We identified two independent variables for volume regulation: cell diameter and the surface-to-mass coupling constant *α*. All other shape variables depend on these two and on the process of cell division. We found that Turgor pressure played an important and potentially regulatory role for cell diameter. In the future, it will be interesting to identify the molecular mechanism underlying the relationship between intracellular mass density and *α*.

## Acknowledgements

We thank Nikolay Ouzounov and Zemer Gitai for strains NO34, NO53, Natasha Ruiz for strain NR693, Suckjoon Jun for plasmid pDB192, Richard Wheeler and Ivo Boneca for help with measuring mDAP incorporation, Waldemar Vollmer for the suggestion to digest sacculi with lysozyme, and Piernicola Spinicelli for technical support on phase microscopy. This work was supported by the European Research Council (ERC) under the Europe Union’s Horizon 2020 research and innovation program [Grant Agreement No. (679980)], the French Government’s Investissement d’Avenir program Laboratoire d’Excellence “Integrative Biology of Emerging Infectious Diseases” (ANR-10-LABX-62-IBEID), the Mairie de Paris “Emergence(s)” program, and the Volkswagen Foundation.

