## Supporting Material and SI Figures for "Bacteria control cell volume by coupling cell-surface expansion to dry-mass growth"

### Materials and Methods

#### 1. Strain and plasmid construction

pBC04: *mepS* was cloned under an arabinose-inducible promoter in the pBAD30 vector. *mepS* was amplified from *E. coli* genomic DNA using the primers 5'-TGACTGACGAGCTCAGGAGGAATTCACCATGGTCAAATCTCAACCGATTTTG-3' containing a Ribosome-binding site (RBS) and a SacI restriction site and 5'-GTCAGTCATCTAGATTAGCTGCGGCTGAGAACCCG-3' containing a XbaI restriction site. After digestion, the *mepS*-containing SacI-XbaI fragment was then ligated with the SacI-XbaI cleaved pBAD30 fragment.

EB06 (MG1655 *lptD4213*) was constructed following (*1*). We transduced MG1655 with P1 phage prepared from NR693 (MC4100 *lptD 4213*, *carB*:Tn10) and selected for growth on carbenicillin and found mutants that did not grow on MacConkey agar. One of the mutants, EB03 (MG1655 *lptD 4213*, *carB*:Tn10), was transduced a second time with the same P1 phage and selected for growth on minimal media supplemented with glucose, which requires *carB* to obtain EB06.

For all other strains, plasmids were transformed using TSS buffer (*2*) and chromosomal changes were performed by P1 phage transduction in the order indicated in Table S1. The kanamycin resistance cassette was excised using plasmid pCP20 (*3*).

#### 2. Growth conditions

Cell cultures were grown from an individual colony in LB Miller at 37C to exponential phase with appropriate selection of antibiotics. Cells were then washed and diluted into LB Miller, defined MOPS minimal medium (MM) (MOPS Minimal Media Kit, M2106, Teknova) or MOPS Rich Defined Medium (RDM) (MOPS EZ Rich Defined Medium, M2105, Teknova). In MM we added mannose (0.4% w/v) (MM+mannose), glycerol (0.4% w/v) (MM+glycerol), glucose (0.2% w/v) (MM+glucose) or glucose (0.1% w/v) and BD Bacto casamino acids (0.1% w/v, Fisher Scientific, 11762764) (MM+glucose+CAA). Osmolality of MM and RDM was measured (Loeser, Freezing Point Osmometer) and adjusted using NaCl to be 280+/-10mOsm.

Before microscopy, we kept cultures in exponential phase for >10 doublings. If not otherwise indicated, cells were grown at 30C in liquid media in a shaking incubator. Strains with plasmids were grown with 50μg/ml carbenicillin. To inhibit division, *sulA* was induced with 1mM IPTG from plasmid pDB192, either during microscopy or shortly before microscopy, as indicated in Table S3.

To strongly repress MepS expression before microscopy, strain b183 (Δ*mepS*, *mreB::msgfp-mreB*)/pBC04 carrying *mepS* under the $P_{BAD}$ promoter was grown in LB Miller with 0.2% glucose in liquid, then washed by centrifugation and resuspension in LB directly before placing cells onto agar pads. To induce MepS expression, we added 0.2% arabinose. In those cells we inhibited cell division using the antibiotic aztreonam (Sigma-Aldrich, A6848, 10 ug/ml), which we included in the agar pad prior to microscopy.

For perturbations of cell-wall synthesis or metabolism, we added the following compounds as indicated in Table S3: Vancomycin (Fisher Scientific, BP2958-1), Fosfomycin (Sigma-Aldrich, P5396-1G), D-cycloserine (Sigma-Aldrich, 30020-1G), alpha-methylglucose (aMG) (Sigma-Aldrich, M9376) with concentrations as indicated in Table S3.

#### 3. Microscopy

Microscopy was carried out on a Nikon Ti-E inverted phase-contrast and epi-fluorescence microscope. Additionally, we added a module allowing for spatial light interference microscopy (SLIM) (see

Supplementary Note for details). The microscope is equipped with a temperature chamber (Stage Top incubator, Okolab) set to 30C if not otherwise indicated. All microscopy was done with a Nikon Plan Apo 100x NA 1.45 Ph3 Objective, which was equipped with a solid-state light source (Spectra X, Lumencor Inc., Beaverton, OR), a multiband dichroic (69002bs, Chroma Technology Corp., Bellows Falls, VT), and excitation (485/25, 560/32) and emission (535/50, 632/60) filters for GFP and FM4-64 imaging, respectively. Epi-fluorescent images were acquired with a sCMOS camera (Orca Flash 4.0, Hamamatsu) with an effective pixel size of 65nm, while phase-contrast and quantitative phase images were obtained with another CMOS camera (DCC3260M, Thorlabs) with an effective pixel size of 87nm.

For SLIM measurements we took six consecutive images with a phase delay of $\frac{n\pi}{2}$, where $n$=1-6, with 200 ms exposure each. Out of these, we obtained three phase images (from images 1-4, 2-5, and 3-6, respectively; Supplemental Note 1) and took the average to obtain the final phase image. Including delays due to software, the acquisition of one final phase image took ~2 s.

For experiments with dividing cells, only individual cells up to their first division were analyzed. For snapshot experiments and for time-lapse experiments of filamenting cells (*sulA* induced), only non-dividing cells were analyzed.

#### 4. Sample preparation

Images were obtained by either immobilizing cells under an agar pad or by immobilizing them in a flow chambers as described below. In all experiments care was taken to image individual cells that are well-separated from any neighboring cells. This is to ensure proper segmentation and integration of optical phase shift. All snapshot images (single-time-point measurements) were conducted on agar pads.  Time-lapse experiments were conducted either on agar pads or in flow chambers, as indicated in Table S3. Agar pads had the advantage to lead to better cell immobilization. However, for cells on agar pads made from minimal media we observed faster than expected growth, demonstrating that the agar pads contain additional nutrients. Time-lapse movies on agar pads were therefore taken only for amino-acid-supplemented media (MM+glucose+CAA or RDM).

Agar pads were prepared from fresh culture medium and 1% UltraPure Agarose (16500-500, Invitrogen). For microscopy, 1ul of cell culture with $OD_{600}$~0.1 was directly applied onto a coverslip (Corning No 1.5), which was already assembled on the microscope and inside the stage top incubator. The droplet was then immediately covered with an agar pad. Optical alignment and focus position had been done before and thus imaging could be started within 0.5-1.5min after taking cells from the shaking incubator. To measure cell dimensions with the membrane stain FM4-64 (ThermoFisher, T13320) for measurement calibration (see

Supplemental Note), we prepared agar pads containing 1% UltraPure agarose, MM+mannose or RDM, and 20ng/ml FM4-64. To avoid fluorescent signal from dirt on the coverslip we pre-cleaned cover slips by bath sonication in a 1M KOH solution for 1h at 40C.

Shifts on agar pads were applied by adding a 1.5ul droplet of medium containing the respective compound (Table S3) to the top of the pad (10x10x1.5mm=150ul) during the time lapse similarly to (*4*), at a time indicated in Table S3. The concentration of the compound in the droplet was adjusted to achieve a final concentration as indicated in Table S3. According to a one-dimensional diffusion model the sample cells experienced 1/10$^{th}$ of the final concentration within approximately 8 min. For the osmotic ramp experiment (Fig. S14), we grew cells in RDM+NaCl (700mOsm), immobilized cells under an 1mm microscopy agar pad (RDM+NaCl, 700mOsm) and eventually placed a 3mm-thick agar pad (RDM, 200mOsm) on top. This leads to a slow decrease in NaCl concentration at the sample with a final osmolality of 325mOsm. According to a one-dimensional diffusion model 90% of the final decrease was achieved after 30min.

For time-lapse experiments in flow chambers we used commercial flow chambers (sticky-slide I Luer 0.2, Ibidi) and provided constant flow of media of 50ul/min via connected silicone tubings using a syringe pump (70-4501, Harvard Apparatus). To attach cells to the surface, 24x60mm coverslips (Corning No 1.5) were pre-treated with APTES ((3-Aminopropyl)triethoxysilane, Sigma-Aldrich, A3648-100ML). Specifically, coverslips were incubated with 2% APTES in EtOH (vol/vol) for 15min at RT, 3x washed in EtOH, rinsed with distilled water and finally stored in EtOH. Before use, coverslips were again rinsed with water and dried with compressed air. Fow chambers were assembled on the microscope and filled with 100ul of cell suspension with $OD_{600}$~0.1. Cells were allowed to settle for ~5min. Shift experiments in flow chambers were conducted by using a Y-junction about 5cm upstream of the channel and switching the flow from syringe #1 (containing medium #1) to syringe #2 (containing medium #2). To avoid leakage of medium #2 into the feeding channel prior to the shift, both inlets of the Y-junction were filled with medium #1 prior to the shift.

#### 5. Immersive refractometry and volume-independent measurement of average refractive index

For immersive refractometry, which allows to determine local cellular refractive index (*5*), we imaged cells using SLIM but increased the refractive index of the surrounding medium. If the cell or part of the cell still has a higher refractive index, it exhibits a positive phase shift. If the refractive index of the medium exceeds the cellular refractive index, the cell shows a negative phase shift. This technique is independent of volume measurements and does not require any correction of the SLIM measurement. For measurements displayed in Fig. S8, MG1655 cells were grown in MM+glucose and in RDM. Additionally, BSA (Sigma-Aldrich, A2153) was dissolved in fresh medium to increase the refractive index to $n_{\text{medium}}$=1.384. Media without and with BSA were adjusted to equal osmolality (~280mOsm). Cells were then immobilized in a flow chamber and the surrounding medium was replaced with MM+glucose+BSA.

Following a different technique to measure the average refractive index of a cell directly (*6*), without using the information about cell shape, we obtained phase images of the same cell for two different media with different refractive indices (Figs. S6, S10; Supplementary Note). To increase refractive index we used high-molecular weight dextran (200kD, 31398, Sigma). Osmolalities of low and high refractive index medium were matched by addition of NaCl (~280mOsm). We exchanged media multiple times, which allowed to measure both the length dependency of dry-mass density between different cells and the decrease of dry-mass density in individual filamenting cells over time.

#### 6. Analysis of shape and mass during time-lapse microscopy

We filtered time traces of mass, length, width, surface, and volume using a Gauss filter of standard deviation σ as indicated in Table S3. For those measurements where we sought to obtain quantities as fast as possible after sample preparation, we used an average filter of width *N* instead (smooth function, MATLAB) (see also Table S3).

Relative rates $\lambda_X = \mathrm{d}(\log X)/\mathrm{d}t$ with $X = A, W, L, M$ with measurement time points $t_i$ were calculated as

$$\lambda_X(t_{i+1/2}) = \frac{2(X_{i+1} - X_i)}{(X_{i+1} + X_i)\Delta t_i},$$

where $t_{i+1/2} = 0.5(t_i + t_{i+1})$. Before display, rates were smoothened with the same filter used for quantities *X*. As a consequence of smoothening, rates seem to change shortly before the timepoint of a shift. For the quantities $\alpha = \mathrm{d}M/\mathrm{d}t$ and $\beta = \lambda_M \alpha$ (Fig. S16), we calculated

$$\alpha(t_{i+1/2}) = \frac{(S_{i+1} - S_i)}{(M_{i+1} - M_i)}$$

and subsequently smoothened with an average filter (smooth function, MATLAB) of width *N* = 9. We obtained $\beta(t_{i+1/2}) = \alpha(t_{i+1/2})\lambda_M(t_{i+1/2})$ and subsequently smoothened with the same filter.

#### 7. Inhibition of cell-wall synthesis and analysis of synthesis arrest

To inhibit cell-wall-precursor production we treated S257 cells grown in RDM with D-cycloserine (*7*, *8*) or Fosfomycin (*9*). To inhibit cross-linking we treated a mutant with permeable outer membrane (*lptD*4213) (*1*, *10*) (S382) with Vancomycin (*11*). Both strains contain an MreB-msfGFP fusion (*12*, *13*) in their native *mreB* loci, which we used to determine the time of MreB-motion arrest. MreB-motion arrest was determined both by visual inspection of movies (Movie S1-3), by kymographs (Fig. 4A), and by the average speed of MreB-msfGFP tracks (Fig. 4A, see also Methods section 8). Once average speed drops below $v$ = 8 nm/s, rotational motion stalled according to visual inspection of movies.

To verify that cells continued to grow after a complete arrest of cell-wall insertion, we complemented our MreB-based measurements with measurements of 3H-mDAP incorporation (Fig. S17, Methods section 9). For efficient labeling and different from the results presented in Fig. 4, we grew cells in MM+glucose+0.4mM lysine. We then determined the time point of cell-wall-synthesis arrest as the time when mDAP incorporation stopped (Fig. S17A). At the same time, we also measured growth by optical density (OD600) (Fig. S17B).

#### 8. Tracking of MreB-msfGFP

MreB-msfGFP images were analyzed using a custom MATLAB code: Images were first filtered in both space and time using a three-dimensional Savitzky-Golay filter with a filter size of 3 pixels in xy-directions time 3 points along the temporal dimension. Images were subsequently de-noised once more using a 2D-Gauss filter ($\sigma$ = 0.5 pixels).

Images were subsequently rescaled by a factor of 5 using spline interpolation to achieve sub-pixel resolution. MreB spots were detected as local maxima inside the cell boundary obtained by segmentation. MreB spots with intensity higher than the cell background is considered for tracking.

The local maxima were connected to construct raw trajectories based on their distance at consecutive time points (*14*) with a maximal displacement during subsequent time frames of 3 pixels. The imaging interval is 2 seconds for a total time of 20 seconds for all movies.

After generating the tracks, we applied a Gauss filter ($\sigma$ = 1.5 pixels) in order to decrease spatial noise. Tracks which have more than 7 localizations are considered for velocity distributions. Velocity is calculated from single displacement vectors of smoothed trajectories.

#### 9. Radioactive 3H-mDAP labelling

S382 (MG1655 *lptD4213 lysA::kanR*/pDB192) was grown in MM+glucose + 0.4mM Lysine following (*15*) and kept below OD600<0.1 for 2 days at 30C ensuring exponential growth for >10 doublings. The culture was split into "cold" culture with 1mM ITPG to inhibit division and "hot" culture with 5uCi/ml 3H-mDAP (MT-1556, Moravek) and 1mM IPTG. After 30min both "hot" and "cold" cultures were split into a control and separate cultures with 100ug/ml Vancomycin, 1mM D-cycloserine and 500ug/ml Fosfomycin, respectively. In regular time intervals, samples of 200ul of each hot culture were taken and boiled in 4% SDS for at least 1h and OD600 readings were taken from each cold culture. After cooling the SDS samples to room temperature, they were filtered through 0.22um membrane filters (Millipore GSWP02500). The filters were then washed with 30ml distilled water, collected into scintillation count vials and treated with 400ul of 10mg/ml Lysozyme at 37C for at least 2h, and finally dissolved in 5ml scintillation cocktail (Filter-Count™, Perkin Elmers) overnight. Eventually, counts per minute due to radioactive decay of 3H were measured in a scintillation counter (TriCarb, Perkin Elmer).

#### 10. Model of mass density of elongating and dividing cells

For non-dividing cells we calculated the modeled mass density $\rho$ using a straight-forward deterministic model of a spherocylinder of fixed width $W$, taken as the average experimental value (Fig. 2F), which increases its surface $S$ in direct proportion to mass $M$, with a ratio $S/M$ given by the average experimental value (Fig. 2E). Due to cell geometry, the surface-to-volume ratio $S/V$ has the length-dependent form

$$\frac{S}{V} = \frac{4}{W\left[1 - \frac{W}{3L}\right]}$$

Dry-mass density can then be written as $\rho = (S/V)/(S/M)$.

To model dividing cells, we consider a cell with surface-to-mass ration $S/M$=16.4 um$^2$/pg, length at birth of $L_b$=2.186 um, doubling time of $t_d$=60 min, and cell-cycle time of initiating the septum of $t_i$ = 42 min (that is, septation takes $t_d$-$t_i$ = 18 min), all determined as averages from the time-lapse experiment displayed in Fig. 2I. For cell width, we used the experimentally determined but time-dependent smoothened average, since width showed weak time dependence (Fig. S11). Septa were modeled as two intersecting hemispheres growing at a rate of constant area increase, as previously demonstrated experimentally (*16*). Here, we adjusted the rate of area increase to match the total duration of septum formation, $t_d$-$t_i$. At the same time, we maintained surface-to-mass ratio constant as observed (Fig. S11).

### Supplementary Tables

**Table S1: Strain list**

| Strain | Genotype | Construction |
|---|---|---|
| MG1655 | WT | Gift from Ghigo lab (Institut Pasteur) (CGSC#6300) |
| NCM3722 | WT | Gift from Rabinowitz lab (Princeton) (*17*) |
| S290 | MG1655/pDB192 | MG1655 → pDB192 |
| NO34 | MG1655 *mreB*::*mreB-msfGFPsw, kanR* | (*12*) |
| NO53 | MG1655 *mreB*::*mreB-msfGFPsw* | (*13*) (difference to NO34: MG1655 is CGSC#6300) |
| S257 | MG1655 *mreB*::*mreB-msfGFPsw*/pDB192 | NO53 → pDB192 |
| EL162 | MG1655 Δ*mepS::kan* | MG1655 → P1(Keio Δ*mepS*) |
| EL54 | MG1655 Δ*mepS* | EL162 → pCP20 |
| b182 | MG1655 Δ*mepS*, *mreB::msgfp-mreB, kanR* | EL54 → P1(NO34) |
| b183 | MG1655 Δ*mepS*, *mreB::msgfp-mreB, kanR*/pBC04 | b182→ pBC04 |
| NR693 | MC4100 *lptD* 4213, *carB*::Tn10 | (*1*) |
| EB03 | MG1655 *lptD* 4213, *carB*::Tn10 | MG1655 → P1(NR693) |
| EB06 | MG1655 *lptD*4213 | EB03 → P1(MG1655) |
| S380 | MG1655 *lptD*4213/pDB192 | EB06 → pDB192 |
| S381 | MG1655 *lptD*4213, *lysA*::*kanR*/pDB192 | SVT380 → P1(Keio Δ*lysA*) |
| S382 | MG1655 *lptD*4213, *mreB*::*mreB-msfGFPsw, kanR*/pDB192 | SVT380 → P1(NO34) |

**Table S2: Plasmid list**

| Plasmid | Genotype | Source |
|---|---|---|
| pCP20 | *FLP*$^{+}$, λ ci857$^{+}$, λ $p_R$ Rep$^{ts}$, Ap$^{R}$, Cm$^{R}$ | (*3*) |
| pBAD30 | Expression vector with $P_{BAD}$ promoter | (*18*) |
| pBC04 | $P_{BAD}$*mepS* | This study |
| pDB192 | *bla* $P_{lac}$::*sulA* | Gift from Jun lab (UCSD) (*19*) |

**Table S3: Microscopy conditions for time-lapse experiments**

Conditions of all time-lapse microscopy experiments including experiment type, corresponding figures, pre-shift (‘Medium #1’) and post-shift media (‘Medium #2’), time point of placing cells on the microscope support ($t_{exp}$) relative to time indicated in figure axis, and microscopy support (agarose pad: ‘pad’, flow chamber: ‘chamber’). If not indicated, *sulA* was induced at $t_{exp}$.
* Medium prior to osmotic shift: MM+glucose+CAA
** At $t$ = 45 min, we added glucose (1 % w/v).
*** $n_{cells}$ increased with time. At $t$ = -30min, $n_{cells}$ > 20
**** At $t_{exp}$ = -15min, *sulA* expression was induced in liquid culture (1mM IPTG)
† for these experiments, σ indicates the width of a smoothing average filter
NA Not applicable

| **Experiment** | **Figure** | **Strain** | **Medium #1** | **Medium #2** | **Δt** (min) | $\boldsymbol{\sigma}$ | $\boldsymbol{t_{exp}}$ (min) | $\boldsymbol{n_{cells}}$ | **support** |
|---|---|---|---|---|---|---|---|---|---|
| Osmotic shock | 1C | S290 | MM* | + NaCl (200mM) | 1 | 0 | -60 | 75 | pad |
| Filamentation | 2G,H; S9 | 290 | MM+glu+CAA | NA | 1 | 25† | 0 | 45 | chamber |
| Division | 2I; S11 | MG1655 | MM+glu+CAA | NA | 1.1 | 0 | NA | 49*** | pad |
| Upshift #1 | 3B; S16 | S290 | MM+mannose | MM+glu+CAA | 1.4 | 1.6 | -101 | 40 | chamber |
| Upshift #2 | 3C; S16 | S290 | MM+glu | RDM | 1.4 | 0.6 | -106 | 40 | chamber |
| Downshift | 3E; S16 | S290 | MM+glu+CAA | MM+mannose | 1.4 | 1.6 | -40 | 25 | chamber |
| correlations | S12 | S290 | RDM | NA | 1.8 | 7† | -27 | 108 | pad |
| correlations | S12 | S290 | RDM | NA | 1.8 | 7† | -27 | 203 | pad |
| Control #1 | S13 | S290 | MM+mannose | NA | 1.4 | 9† | -9 | 35 | chamber |
| Control #2 | S13 | S290 | MM+glu | NA | 1.2 | 9† | -18 | 130 | chamber |
| Control #3 | S13 | S290 | MM+glu+CAA | NA | 1.4 | 9† | -11 | 26 | chamber |
| aMG | S15 | S257 | MM+glu (0.01% w/v) | +aMG (0.5 % w/v) | 1 | 0.6 | -40 | 80 | pad |
| Mecillinam | 3F | S257 | RDM | +mec (100 ug/ml) | 1 | 1 | -40 | 15 | pad |
| Vancomycin | 4B | S382 | RDM | +van (100 ug/ml) | 1.1 | 1 | -20**** | 37 | pad |
| D-cycloserine | 4B | 257 | RDM | +dcyc (1 mM) | 1.1 | 1 | -22**** | 60 | pad |
| Fosfomycin | 4B | 257 | RDM | +fosf (500 ug/ml) | 1.1 | 1 | -24**** | 81 | pad |
| Van+aMG | 4C | S382 | MM+glucose (0.02% w/v) | +aMG (0.5 % w/v)** | 1 | 1 | -58 | 77 | pad |
| Hypo-osmotic ramp | S14 | S290 | RDM+NaCl (702mOsm) | RDM (325mOsm) | 1 | 0 | -24 | 27 | pad |
| Osmotic ramp control | S14 | S290 | RDM+NaCl (697mOsm) | RDM (325mOsm) | 1 | 0 | -24 | 26 | pad |
| MepS | 4D | b183 | LB Miller | + arabinose (0.2% w/v) | 1.4 | 1 | -26 | 11 | pad |
| Immersive refractometry | S6,S10 | S290 | MM+glu+CAA (280mOsm) | MM+glu+CAA+ Dextran (20% w/v) | 0.3 | 0 | -15 | 29 | pad |
| Fixed cell | S5 | S290 | RDM | NA | 0.04 | 0 | NA | 1 | pad |

### Supplementary Note – Quantitative Phase Microscopy

#### 1. Microscope setup

For dry-mass measurements we used Spatial Light Interference Microscopy (*20*), which we added to a Nikon Ti-E microscope using custom components (Fig. S1). The Nikon Ti-E is equipped with typical phase-contrast illumination with illumination confined by an annulus (Ph3) in the condenser (ELWD NA 0.52) and a matching phase-ring in the pupil plane of the objective (Plan Apo 100x 1.4 Ph3). To fully illuminate the condenser aperture, we added a 2x telescope to the illumination arm of the Ti-E (LC1315-A and LA1417, Thorlabs). The pupil plane of the objective is relayed via a fourier lens (AC508-300-A, Thorlabs) onto a spatial light modulator (SLM, Meadowlarks P1920) where a matching phase ring is displayed. This allows to modulate the phase ring (see below for details). Another fourier lens (AC508-200-A, Thorlabs) completes the first optial 4f-configuration. Another 4f-configuration (two AC508-200-A lenses, Thorlabs) is used to access the pupil plane again, where an iris aperture (Thorlabs) limits the objective numerical aperture to N.A.~0.6. Finally, a set of long and short pass filters (FEL0600 and FES0700, Thorlabs) restricts light transmission to 600-700nm, which in combination with the power spectrum of the illumination LED gives a center wavelength of ~635nm. The image plane is then projected onto the camera. Finally, a linear polarizer (LPVISE200-A, Thorlabs) is placed near the image plane at the exit port of the microscope and orientated with the SLM to act in “phase modulation” mode.

The microscope and display of the SLM monitor were controlled via MATLAB. Micro-manager was used to control the Nikon Ti-E microscope and acquire images from the camera from within MATLAB.

#### 2. Spatial Light modulator calibration and phase modulation

The SLM is connected via HDMI to a PC and acts like a second monitor such that the image displayed on the second monitor is fed to the SLM and the gray-value signal of each pixel on the monitor controls the corresponding pixel on the SLM. The phase delay imparted by the SLM was calibrated as described in (*20*). Briefly, a linear polarizer was placed 45° to the active axis of the SLM, such that it works in “amplitude modulation” mode. The transmitted intensity during a gray-value sweep was recorded and the phase delay was obtained via a Hilbert transform of the amplitude response:

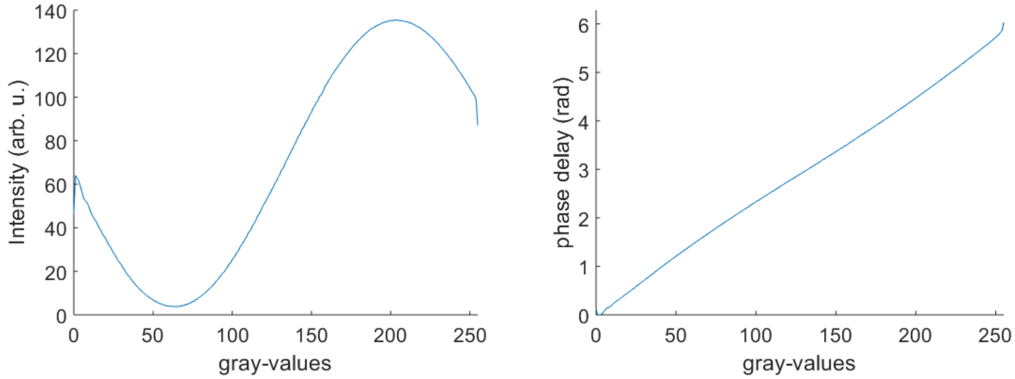

***SLM calibration:*** *Intensity response with SLM in amplitude modulation mode (left). Conversion of gray-values into phase delay (right)*

Furthermore, we confirmed a sufficiently long coherence length of the illuminating white light by retrieving the autocorrelation from its optical spectrum, giving ${l_c}^{\mathrm{FWHM}} = 4.7$ um and several full cycle modulations in the flat part near the central peak. This allows the application of the phase shifting procedure in SLIM (*21*, *22*).

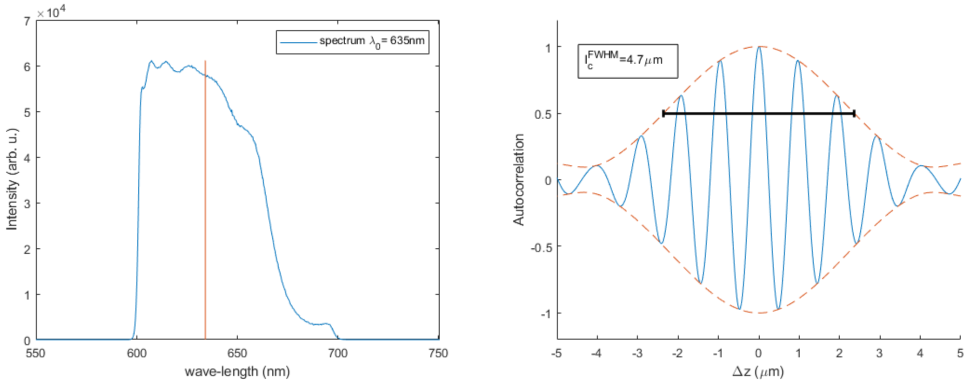

***Illumination Spectrum (left) and Autocorrelation (right):*** *White-light illumination source behaves like a monochromatic field with wavelength λ=635nm and a coherence length of less than 5um*

#### 3. Optical phase retrieval

The optical phase shift of light travelling through the cell can be obtained from 4 intensity images as outlined in (*20*, *22–24*). Conceptually, the total optical field $U_t$ emanating from the sample can be decomposed into the incident field $U_i$ and scattered field $U_s$ (*24*). In phase-contrast microscopy, these fields are mainly spatially separated in the pupil plane of the objective: $U_i$ passes through the phase ring, while $U_s$ mainly passes outside the phase ring. Due to the presence of the phase-ring in the objective and again on the SLM, the incident field can be phase-shifted with respect the scattered field. For SLIM this is done in increments of $\pi/2$ such that the total field becomes:

$$U_t(x,y;n) = U_i e^{i\frac{n\pi}{2}} + U_s(x,y).$$

Here, $n$ is the number of π/2 increments due to phase ring and SLM ring. The image $I_n$ on the detector is the result of the interference of $U_i e^{i\frac{n\pi}{2}}$ and $U_s$:

$$I_n(x,y) = |U_i e^{i\frac{n\pi}{2}} + U_s(x,y)|^2 = |U_i|^2 + |U_s(x,y)|^2 + 2|U_i||U_s(x,y)|\cos\left(\Delta\varphi(x,y) + \frac{n\pi}{2}\right),$$

where $\Delta\varphi(z,y) = \varphi_i - \varphi_s(x,y)$. $I_n(x,y)$ is obtained directly from the camera images after correcting for background signal. The aim of SLIM is to retrieve the optical phase shift of the cell with respect to the incident light, $\varphi = \varphi_i - \varphi_t$. $\varphi$ is obtained as

$$\varphi = \arg\left(\frac{\beta\sin(\Delta\varphi)}{1 + \beta\cos(\Delta\varphi)}\right),$$

where $\Delta\varphi$ is the phase shift between $U_i$ and $U_s$ and $\beta$ is the ratio of field amplitudes $|U_s|/|U_i|$. Following ref. (*23*), $\Delta\varphi$, in turn, is obtained as

$$\Delta\varphi = \frac{I_3 - I_1}{I_4 - I_2}$$

And $\beta$ is obtained as

$$\beta = \gamma\,\frac{1}{4I_i}\frac{I_4 - I_2 + I_3 - I_1}{\sin(\Delta\varphi) + \cos(\Delta\varphi)}.$$

Here, $\gamma$ accounts for the amplitude attenuation of the phase ring in the objective. In the case of the objective used here (Nikon 100x 1.45 Ph3), we imaged the backfocal plane and measured the amplitude attenuation to be $\gamma = \sqrt{0.1}$. The illumination intensity $I_i$ in the expression for $\beta$ is obtained as

$$I_i = \frac{g + \sqrt{g^2 - 4L^2}}{2}.$$

Here, $g$ and $L$ are defined as

$$g = \frac{I_1 + I_2 + I_3 + I_4}{4},$$

$$L = \frac{1}{4}\frac{I_4 + I_3 - I_2 - I_1}{\sin(\Delta\varphi) + \cos(\Delta\varphi)}.$$

$I_i$ should be homogenous across the field of view, but the estimate $I_i$ tends to be biased near the presence of phase objects. To avoid bias on the estimate of $I_i$ at or near a cell, the pixels within ~4 um distance of any phase object are excluded and $I_i$ is inferred from the surrounding area of every cell. Specifically, the intensity field $I_i$ is smoothened before exclusion of phase objects. Then, the intensity values inside regions of phase objects is interpolated. The process of phase retrieval is also illustrated in Fig. S2.

#### 4. Increased focal depth to measure full phase delay

The optical setup contains an iris aperture to limit the NA of the objective (position in the second 4f-configuration, conjugate to the pupil plane). Thus the focal depth is increased and the phase signal can be obtained for the entire cell. Below the integrated phase signal of a 370nm polystyrene bead with increasing defocus dz is plotted. Given the small variation of <1% for |dz|<0.4um and taking into account the dimension of the bead, this indicates that the total phase shift imparted by a cell can be measured for cells with a diameter of up to ~1.2um. Larger defocusing indicates that even the phase shift of an object of ~2.4um diameter would still be measured to more than >90%.

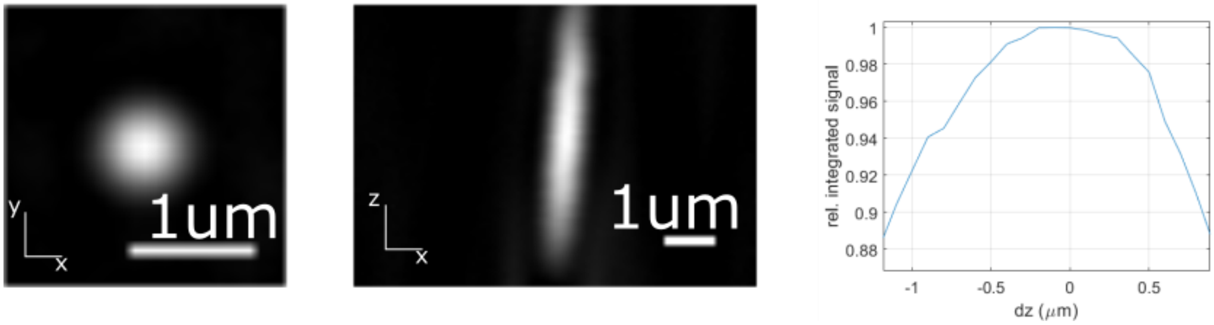

*Left: Quantitative phase image of 370nm Polystyrene bead with the objective's effective NA limited to ~0.6. Middle: z-projection clearly showing an extended depth of focus. Right: Normalized integrated phase of the bead along z*

#### 5. Optical path-length noise level

SLIM has been shown to exhibit a low background noise level (*20*). To demonstrate this we measured the phase signal inside a flow chamber filled with water, but 100um away from the coverslip. This should demonstrate the intrinsic noise level associated with our setup irrespective of the biological sample. We find a uniform background across nearly the entire field-of-view with a standard deviation in the measured optical path-length of std = 0.001 rad. This is <0.01% of the integrated phase shift of a typical cell.

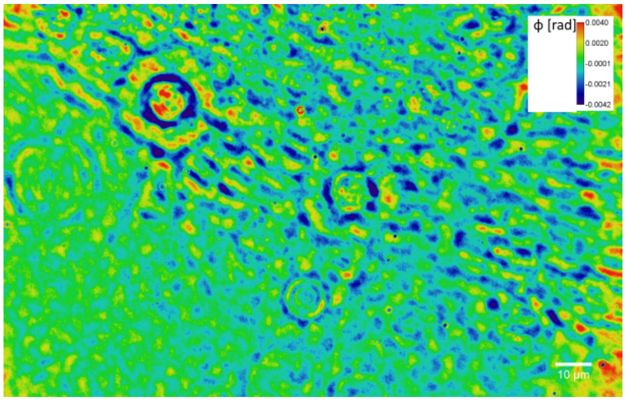

*Optical path-length background noise: Pseudo-colored between +/-0.004*rad*. Scale bar 10 um.*

#### 6. Cell shape and volume measurements and corrections

Cell dimensions were obtained from phase-contrast images acquired using the SLIM module (Fig. S3). We used the MATLAB based tool Morphometrics (https://simtk.org/projects/morphometrics), (*25*) to determine cell contours. The image-formation process through the microscope, but also the contour-finding routines of Morphometrics can bias and distort the contour. We correct and calibrate for this as follows: We base our calibration on images of cells stained with the fluorescent membrane stain FM4-64. These images were acquired using the epi-fluorescent port of the microscope. To correct for diffraction, we also simulated membrane stained cells using the MATLAB based tool BlurLab (https://simtk.org/projects/blurlab/), (*25*). To this end we acquired the point-spread function (PSF) from 100nm fuorescent beads (TetraSpeck, Thermo Fisher) and used BlurLab to convolve point-emitters sitting on the surface of a sphero-cylinder, with the PSF. We applied the correction found in silico onto the measured membrane contours to obtain an estimate for the true contour of the periphery of the cell. In addition to the epi-fluorescent images, we obtained phase-contrast images of the same cells using the SLIM module. We then overlaid the measured contour of the phase-contrast cell with the corrected fluorescent membrane contour and found the radial offset between the two (grey lines in right panel of Fig. S3). Doing this comparison for $n$>1000 cells grown in MM+mannose and in RDM, we inferred an average correction as a function of distance from the cell pole. We found no significant dependence on cell width and growth medium and therefore only applied corrections as a function of distance from the pole. This correction was used to correct the contours of all cells measured with the SLIM module.

Finally, given the calibrated contours of the cell, we use Morphometrics to apply a mesh-grid of 1px (87nm) step-size. This routine also gives the centerline of the cell, which is used to determine cell length. We then assume cylindrical symmetry around the centerline and infer cell surface and cell volume from the sum of the surfaces and volumes of truncated conical wedges with height and width given by the meshes (Fig 1B).

#### 7. Correction of phase signal via comparison with simulated images

The estimate of the optical phase shift outlined in section 3 assumes complete separation of incident field and scattered field at the Fourier plane (i.e. in the plane of the objective phase ring and the plane of the SLM). However, as a consequence of the finite thickness of the phase ring, low frequency components of the scattered field leak into the ring and are incorrectly phase-modulated. For the same reason phase-contrast images are typically plagued with halos and shade-off artifacts. Thus depending on the shape of the phase object, the estimate $\varphi$ is under-estimated, with larger objects being more affected than smaller/finer structures (*24*).

In order to account for the attenuation of the measured phase of a cell, we simulate images of phase objects of the same shape and dimensions as the cell (Fig. S4). The image formation in phase-contrast microscopy (*26*, *27*) and in SLIM (*24*) with partially coherent illumination is well described and we use adaptations of available software packages to simulate images (https://github.com/mehta-lab/microlith; https://github.com/thnguyn2/Halo_removal_with_two_gammas). In essence, the partially coherent image formation can be described as the summation of images illuminated from coherent point sources. We could measure all optical parameters, like size and thickness of the phase ring, illumination ring (location of point sources) and attenuation of the phase ring directly by imaging the pupil plane of the objective. Thus, it is possible to simulate the image formation from an input ground-truth to the measured image without any free parameters. The input ground-truth was modelled as a phase-only object of homogenous density with dimensions found from the corrected cell contour of phase-contrast image. The density was adjusted such that the average phase of the simulated cell matches the measured phase of the actual cell inside the cell boundaries determined from the phase-contrast image. We then obtain a correction factor as the ratio of the integrated phase of input over output. The measured integrated phase of each cell (obtained by summing over all phase-positive pixel values both inside and outside the cell contour) is then multiplied with the correction factor to obtain the corrected integrated phase.

To decrease computational cost of the simulation, we limited the number of point sources on the illumination ring to 75 point sources evenly distributed across the illumination ring. However, we verified that increasing the number of points did not change the phase signal of the simulated cell. This simulation is done for every cell at every time point in all SLIM experiments.

#### 8. Conversion of phase shift to cellular dry mass

The concept to relate refractive index to the dry-mass density of a cell has been proposed by Barer (*28*). This idea relies on the assumption that the refractive index of a cell is proportional to its dry matter content:

$$n_{\text{cell}} = n_{\text{H2O}} + \gamma\rho \,,$$

where $\rho = M_{\text{dry}}/V_{\text{cell}}$ is the average concentration of dry matter of the cell. $\gamma$ is a proportionality constant called the refraction increment, which reflects the average composition of the cell. The linear relation between refractive index and mass density has been shown for well-mixed solutions of different solutes over a wide range of concentrations (*29*, *30*). According to our own osmotic-shock experiment (Fig. 1B) and to similar experiments in HeLa cells (*31*), a linear relationship between refractive index and mass density apparently also holds for the complex interior of a cell. We use an average refraction increment of $\gamma =$ 0.175 ml/g, which we obtained by considering the reported composition of dry-mass (*32*) and reported values for the refraction increments (*5*, *30*, *33*–*36*):

| Refractive increment (ml/g): | % of total dry weight |
|---|---|
| DNA = 0.17 | 3.1 |
| RNA = ~0.18 | 20.4 |
| Proteins=0.185 | 55.0 |
| Lipopolysaccharide 0.15 | 3.4 |
| Phospholipids=0.16 | 9.1 |
| Glycogen=0.15 | 2.5 |
| Cell wall=~ 0.18 | 2.5 |
| Metabolites | 2.9 |
| Ions (e.g. KCl=0.12; NaCl=0.16) | 1.0 |

We also took into account a ~1% decrease of the refraction increment due to wavelength and temperature (*29*).

The dry-mass composition reported by Neidhardt (*32*) describes the *Escherichia coli* B/r strain grown in MM+glucose. Between fast and slow growth, composition changes (*37*). However, the biggest change in the cellular composition is an increase of RNA and decrease of protein relative to the total dry mass (*37*). Since RNA and protein have similar refraction increments, the average refraction increment stays nearly constant across growth rates. We therefore assume a constant refraction increment throughout. We note that the absolute accuracy $\gamma = 0.175$ ml/g is not important for any of our conclusion, which are all based on relative changes of dry mass.

Dry mass is then obtained as the area integral of the optical phase plus a correction due to the refractive index of the medium, $n_{\mathrm{medium}}$, which is higher than the refractive index of water, $n_{\mathrm{H2O}}$:

$$M_{\mathrm{dry}} = \frac{1}{\gamma}\left[\frac{\lambda}{2\pi}\int \varphi \mathrm{d}x\mathrm{d}y + V_{cell}(n_{\mathrm{medium}} - n_{\mathrm{H2O}})\right].$$

Here, λ is the central wavelength. For our experiments, we measured refractive index $n_{\mathrm{medium}}$ at 20C using a refractometer (Brix/RI-Chek, Reichert). We then corrected measured values for our microscope temperature of 30C.

#### 9. Precision of shape and mass measurement of fixed cell

To assess the precision of our shape and dry-mass measurements, we fixed MG1655 cells grown in RDM by treating with 3.7% formaldehyde for 20 min at room temperature. After washing with PBS we imaged cells on an agar pad (1% Ultrapure agarose in PBS). Single-cell precision of shape and mass measurements show a coefficient of variation of CV<0.2%, with dry-mass measurements showing CV=0.06% (Fig. S5). This indicates that our implementation of SLIM and subsequent processing to dry mass surpasses the implementation of SLIM reported by Mir et al. (*38*).

#### 10. Exponential growth in dry mass with correction factor from simulation

The importance of the correction factor derived from simulated images (section 7) can nicely be seen following the growth dynamics of exponentially growing bacteria. Cells grow exponentially in mass (*39*). We also see exponential growth in dry mass (Fig. 2G), but only after correction from the simulation. Without correction the integrated phase does not show constant exponential growth due to the shape-dependent correction factor.

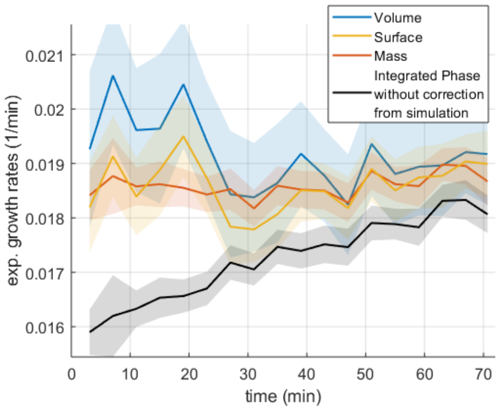

*Constant exponential growth after correction from simulation. Cells were imaged on an RDM agar pad. (solid lines: avg; shading: SEM; <L(t=0)>=4.6um; <L(t=75min)>=18.5um)*

#### 11. Calculation of average refractive index based on media with different indices

Comparison of the integrated phase shift $\varphi_1$ and $\varphi_2$ obtained from the same cell, but in two different media of different refractive indices allows to deduce the average refractive index $n_{\text{cell}}$ as follows (*6*):

$$\varphi_1 = \frac{2\pi}{\lambda}(n_{\text{cell}} - n_{\text{medium}})V_{\text{cell}}$$

$$\varphi_2 = \frac{2\pi}{\lambda}(n_{\text{cell}} - n_{\text{medium}} + \delta_{\text{Dextran}})V_{\text{cell}}$$

$$n_{\text{cell}} = \frac{\delta_{\text{Dextran}}\varphi_1}{\varphi_1 - \varphi_2} + n_{\text{medium}}$$

Here, $\delta_{\text{Dextran}}$ is the refractive index difference due to dextran and $n_{\text{medium}}$ is the refractive index of the medium without dextran. We convert $n_{\text{cell}}$ to dry-mass density using a refraction increment of $\gamma = 0.175\ \text{ml/g}$.

### Supplementary figures

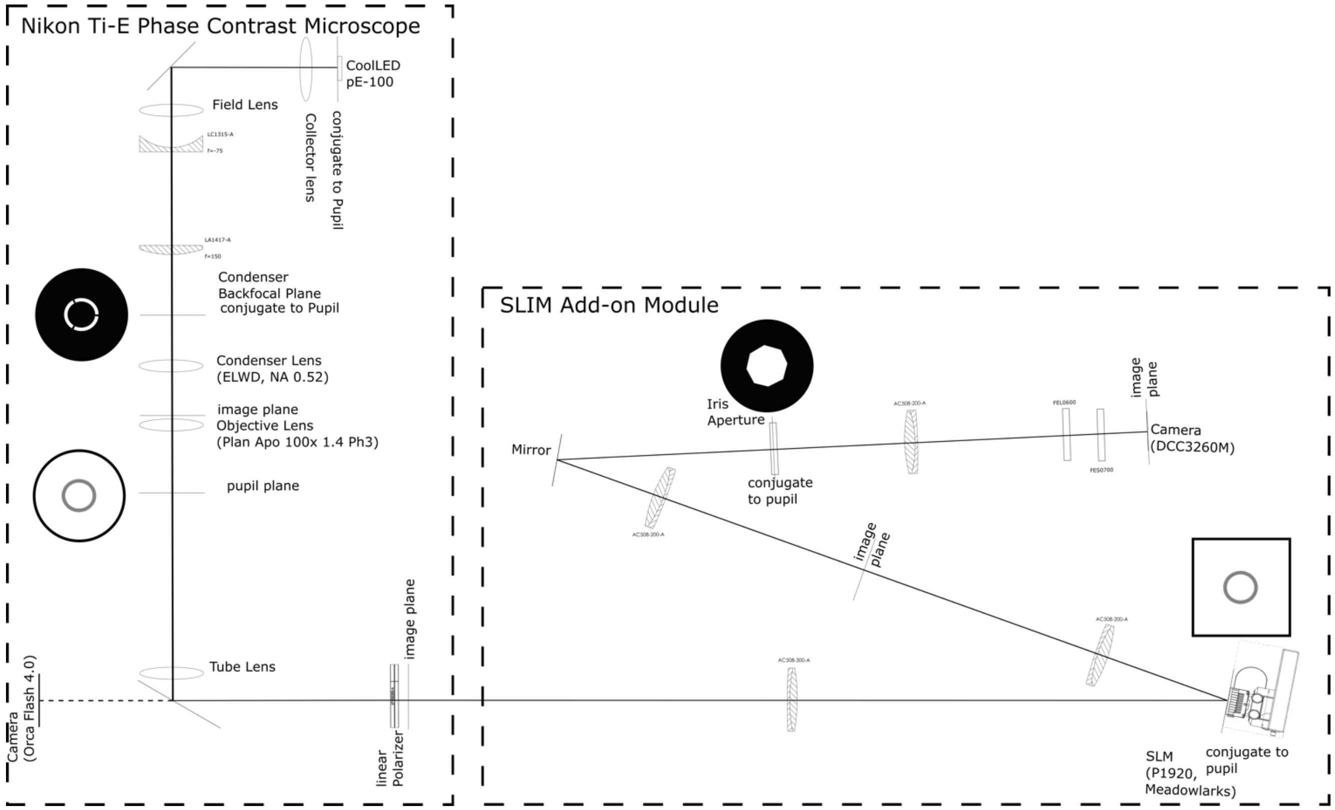

**Fig. S1. Microscopy setup.** Thick dashed boxes indicate the Nikon microscope and the SLIM add-on module. Solid black line indicates optical axis.

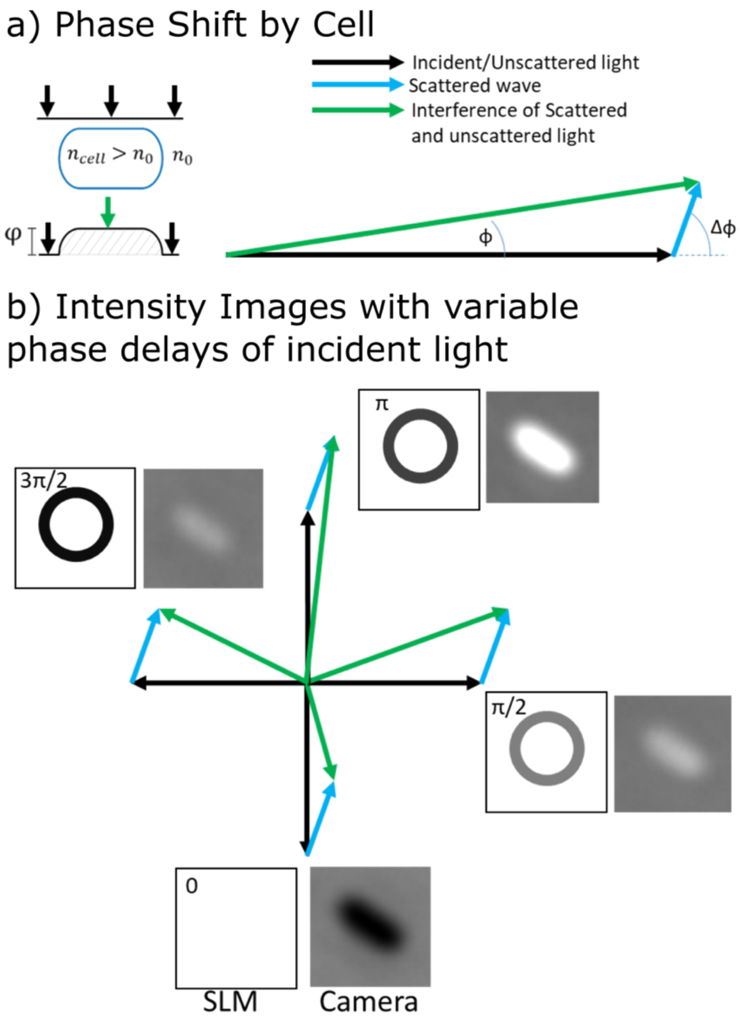

**Figure S2. Optical phase shift and intensity-image formation in SLIM. A:** Left: A cell with refractive index higher than the surrounding $n_{\text{cell}}>n_0$ gives rise to a positive optical phase shift φ. Right: Illustration of resulting wave-vectors. **B:** Analogous to phase-contrast microscopy, the display of the ring on the SLM matching the ring in the objective allows to impart phase delays onto the unscattered light, while the scattered light remains nearly unaffected. Four intensity images with phase delays of $\lambda/4$ increments carry sufficient information to reconstruct the optical phase shift of the light travelling through the cell.

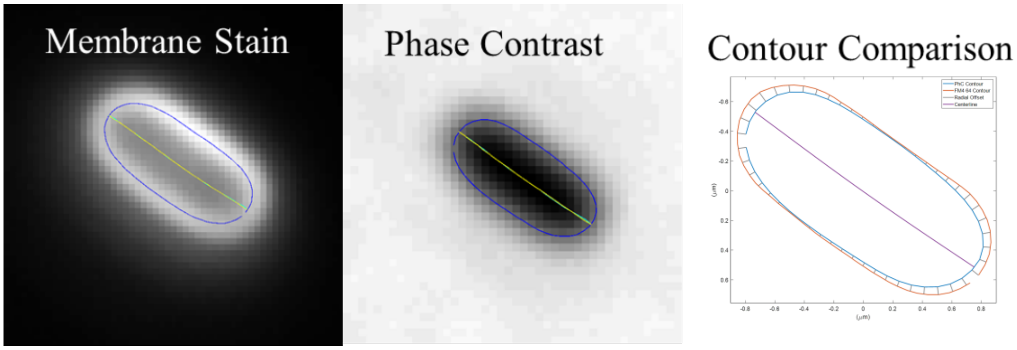

**Fig. S3. Contour calibration.** Cell contour found from FM4-64 membrane stain (left) and cell contour from phase contrast image (middle) is overlaid (right). Radial offset between contours (grey lines) is measured a function of distance from the pole. Average radial offsets from *n*>1000 cells are used to correct contours in all experiments.

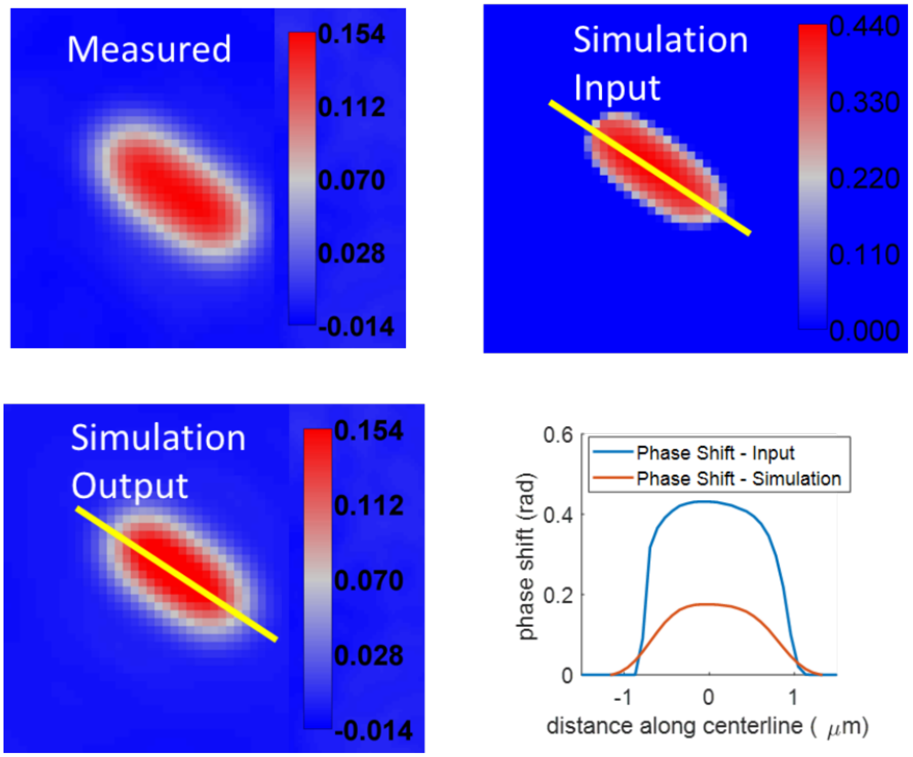

**Fig. S4. Phase-shift correction.** The measured phase image (top left) suffers from optical artifacts. Cell shape (from phase-contrast image according to Fig. S3) is used to create a phase object (simulation input, top right) to simulate the quantitative phase image (simulation output, bottom left). Color scale indicates phase shift. The ratio of input and simulated output phase shift (bottom right) within cell boundaries is used as correction factor for the calculation of the integrated phase shift from the measured phase image.

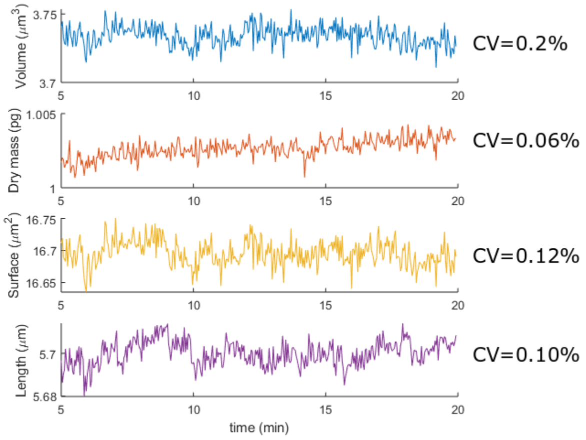

**Fig. S5. Precision of shape and mass measurements.** Recordings of volume, dry mass, surface, length from a single fixed cell previously grown in RDM. Coefficients of variation along time are shown on the right.

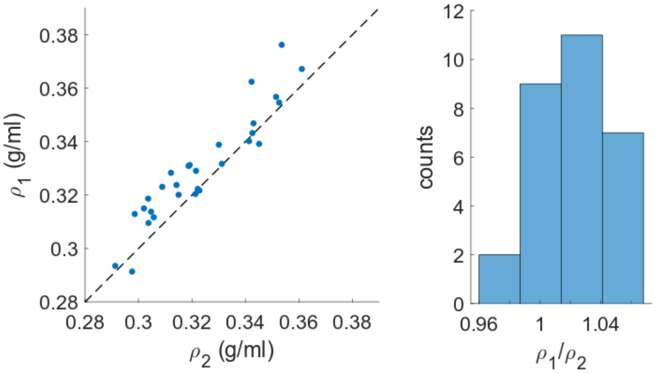

**Fig. S6. Alternative measurement of mass density through immersive refractometry.** We determined mass density of filamenting cells (S290) growing in MM+Glucose+CAA in a flow chamber in two different ways: In Method 1 we calculate volume based on cell contour and mass based on a single phase image plus simulation-based correction, as described in the main text and in Supplementary note sections 6-7. In Method 2, referred to as immersive refractometry in the main text, we took two phase images of the same cells before and after changing the refractive index of the medium. We then measured the average refractive index of the cell directly (Supplementary Note section 11). Both methods give similar results (left panel) with a ratio $\langle \rho_1/\rho_2 \rangle = 1.021 \pm 0.023$ (std). The deterministic offset of 2.1% is within the uncertainty of method 2, due to the uncertainty in measuring the difference in refractive index between the two media. Notably, method 2 does not require an independent volume measurement, nor a correction of the phase signal through simulation. Thus, method 2 validates volume measurement and phase correction in method 1.

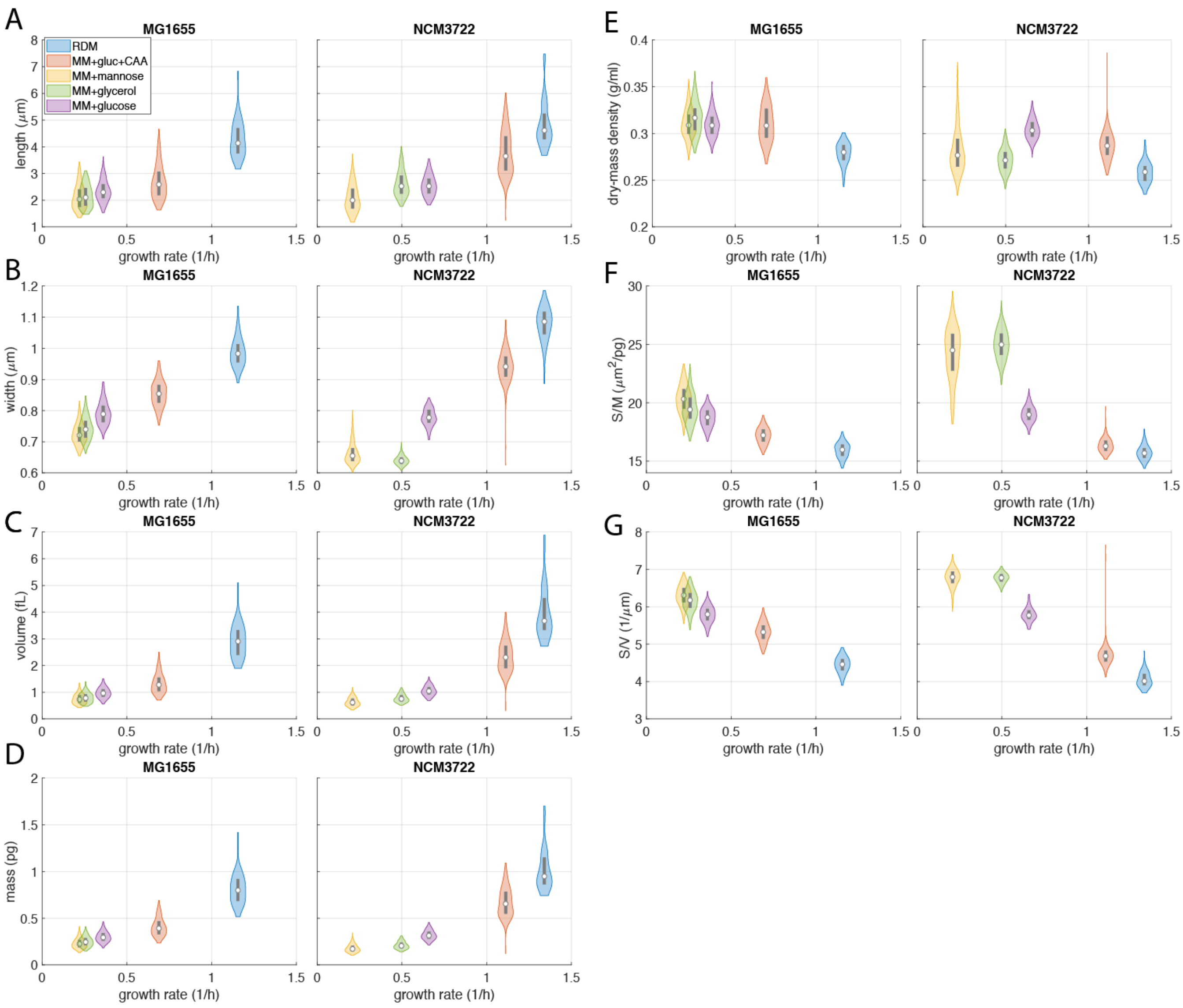

**Fig S7. Single-cell properties of MG1655 and NCM3722 during steady-state exponential growth in different nutrient environments.** Corresponds to Fig. 2B-F. **A:** Cell length, **B:** Width, **C:** Volume, **D:** Mass, **E:** Dry-mass density, **F:** Surface-to-mass ratio, **G:** Surface-to-volume ratio. (white circles=median; grey rectangles=interquartile range)

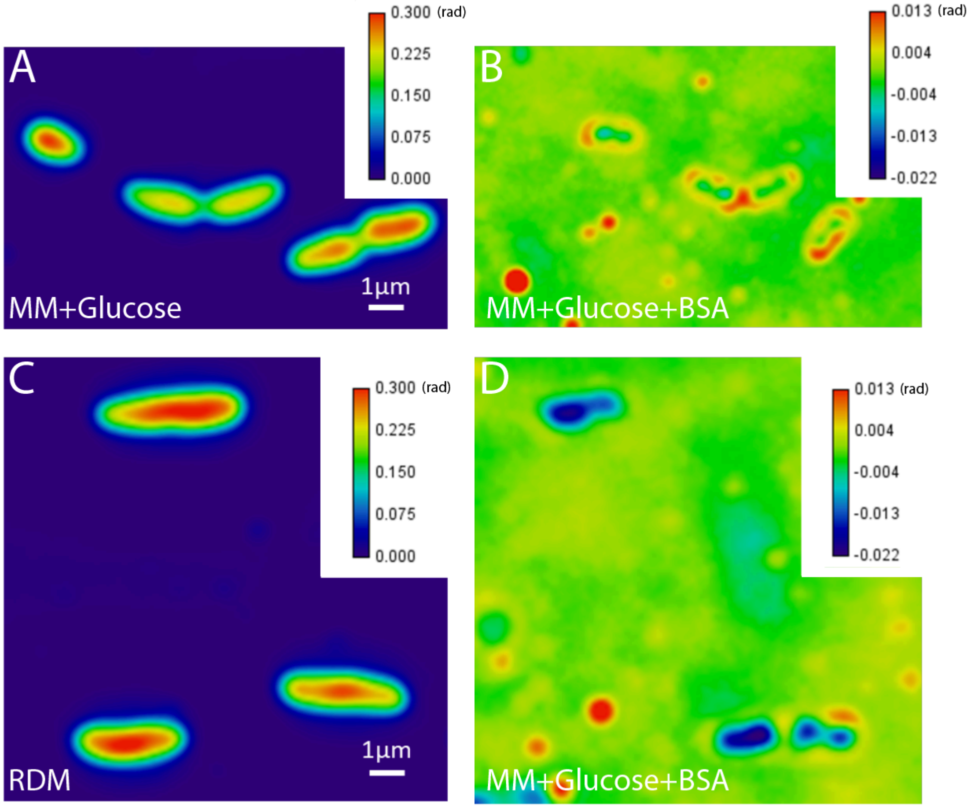

**Fig S8. Immersive refractometry confirms systematic variations in dry-mass density between different media:** SLIM images of WT cells (MG1655) grown in either MM+glucose (**A,B**) or RDM (**C,D**) in a flow chamber, before (left) and after (right) exchange to MM+glucose+BSA ($n_{medium}$=1.384, which corresponds to a dry-mass density of ~0.29g/ml). Note that $\varphi<0$, if $n_{cell}<n_{medium}$. In MM+glucose-grown cells, phase shift is positive ($n_{cell}$>1.384), in RDM-grown cells, phase shifts is negative ($n_{cell}$<1.384). This confirms our measurements based on integrated phase and volume inferred from cell shape (Fig. 2B).

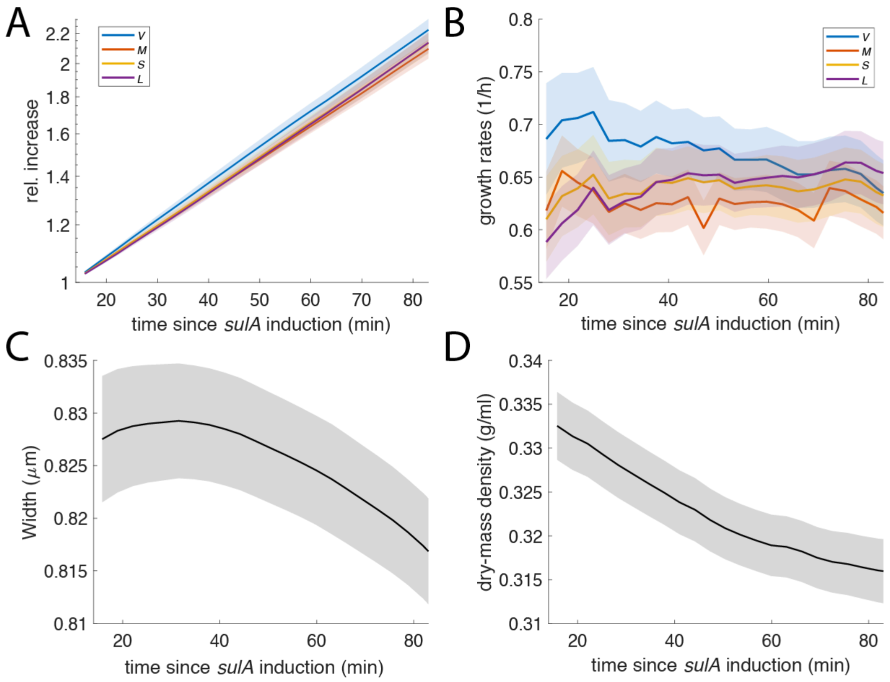

**Fig S9. Single-cell time lapse of filamenting cells.** Single-cell time lapse of filamenting cells (S290) in flow chambers (same experiment shown in Fig. 2G,H). **A, B:** Relative increase (A) and single-cell rates (B) of volume, mass, surface, and length. **C:** Width. **D:** Dry-mass density.

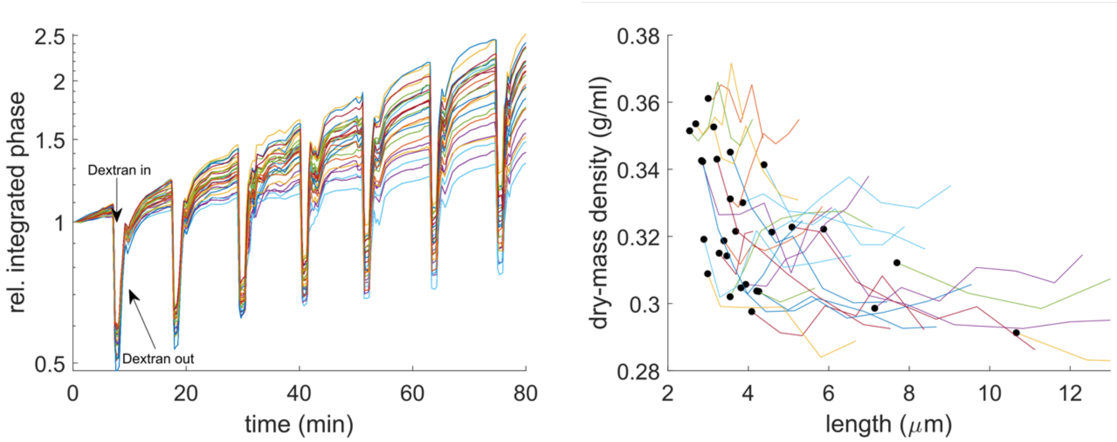

**Fig. S10. Immersive refractometry confirms length dependency of dry-mass density.** Filamenting cells (S290) growing in MM+Glucose+CAA in flow chamber. MM+Glucose+CAA+Dextran with equal osmolality was repeatedly flushed in and out leading to changes in refractive index of the medium and thus changes in integrated phase (**left**). **Right:** Length dependency of dry-mass density. Dry-mass density was inferred from integral refractive index and was measured repeatedly over the course of one doubling. (black dots=first time point; colored lines=single cell traces)

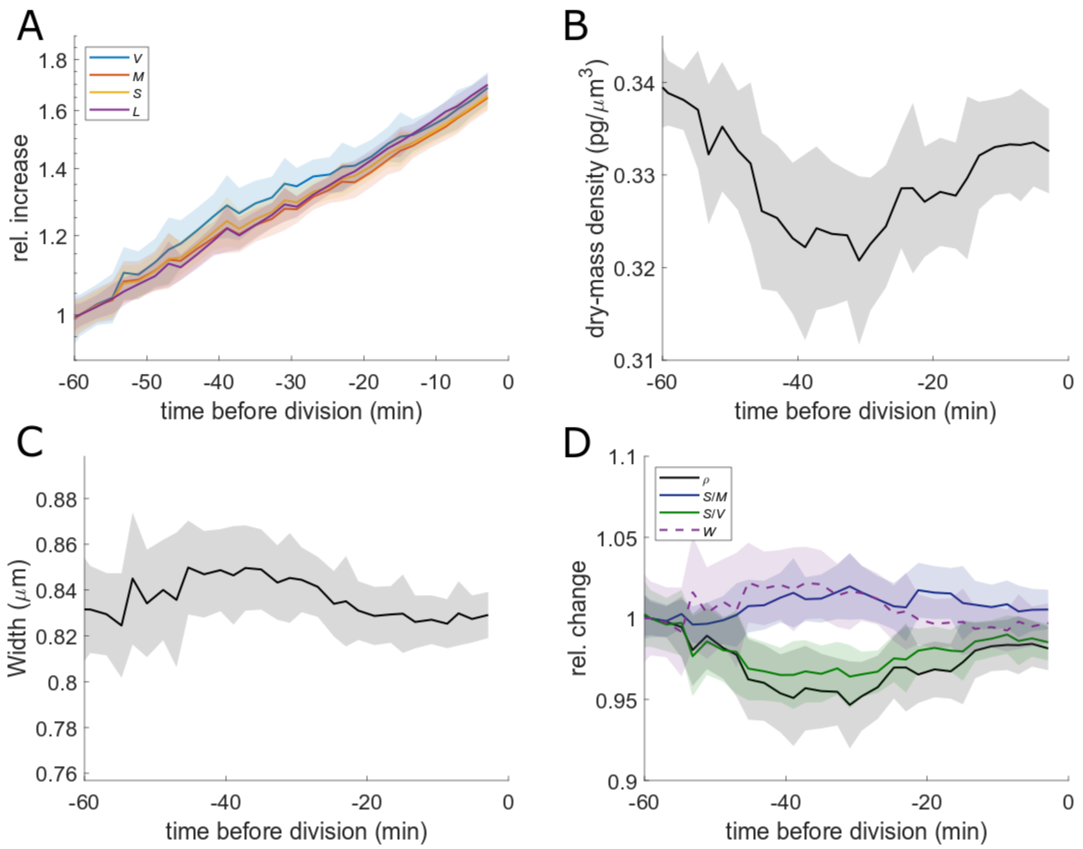

**Fig S11. Dividing cells:** Single-cell time lapse of dividing cells (MG1655) growing on agar pad with MM+glucose+CAA. Single cell traces were aligned in time with respect to their first division, such that $t$ = 0 min is time of division. Relative increase (**A**) of volume, mass, surface, and length. Dry-mass density (**B**), width (**C**), and relative changes of dry-mass density, surface-to-mass- and surface-to-volume ratios, width (**D**). (solid lines+shadings= average±2*S.E.M.)

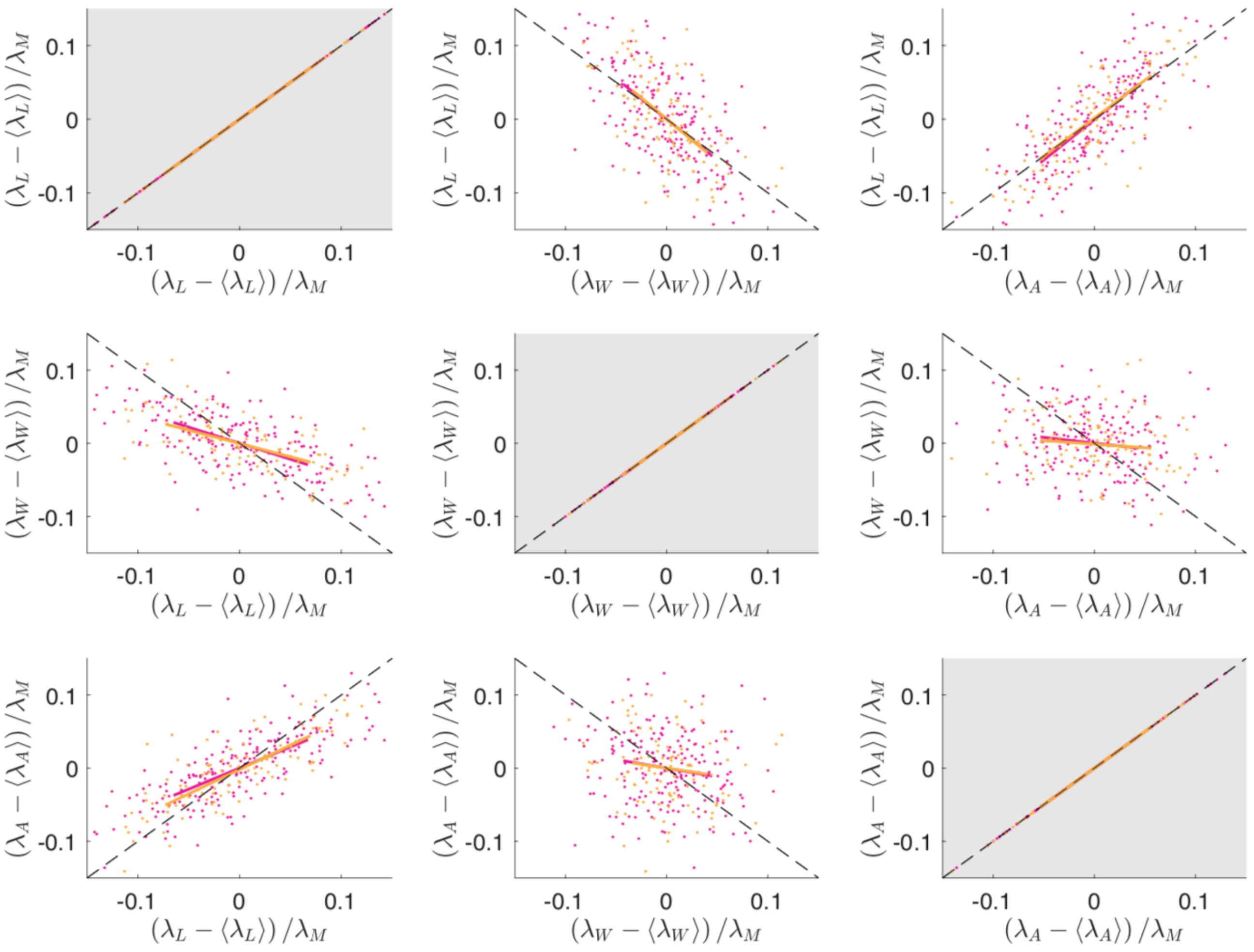

**Fig. S12. Single-cell correlations support the dependency of elongation rate on surface and widening rates.** Matrix of instantaneous correlations between normalized single-cell rates measured from filamenting cells (S290) growing in RDM on agar pad. Exponential rates are defined as $\lambda_{A,W,L,M} = \mathrm{d}(\log A, W, L, M)/\mathrm{d}t$. Two colors represent two identical replicates. Lines: linear regression. $\lambda_L$ shows strong correlations with $\lambda_W$ and $\lambda_A$ (top row), while $\lambda_W$ (middle row) and $\lambda_A$ (bottom row) show weaker dependency on any of the three quantities, suggesting that $\lambda_L$ is a function of $\lambda_A$ and $\lambda_W$, while $\lambda_A$ and $\lambda_W$ are independently coupled to mass growth (Figs. 2,3) and Turgor (Fig. 3,S14), respectively.

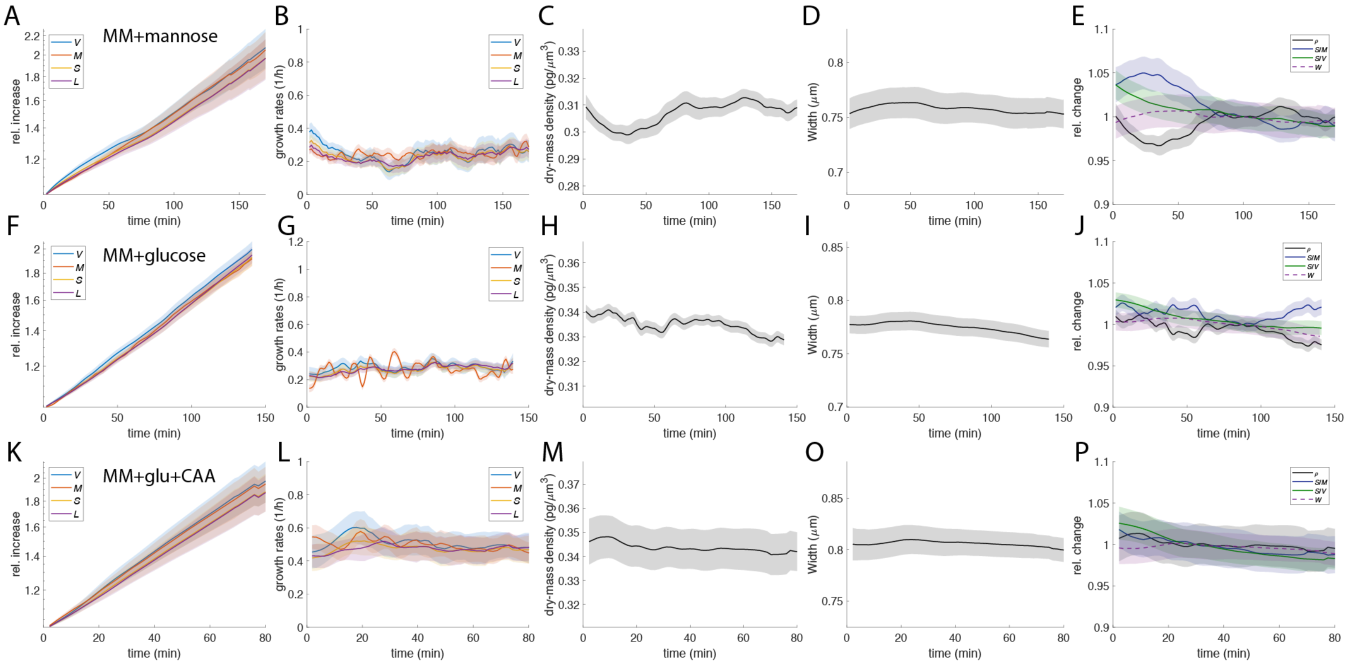

**Fig. S13. Control experiments for all time-lapse nutrient shifts.** Single-cell time lapses of filamenting cells (S290) growing in flow chambers with MM+mannose (top), MM+glucose (middle), and MM+glucose+CAA (bottom). From left to right: Relative increase (A,F,K) and single-cell rates (B,G,L) of volume, mass, surface, and length. Dry-mass density (C,H,M), width (D,I,O), and relative changes of dry-mass density, surface-to-mass- and surface-to-volume ratios, width (E,J,P). (solid lines+shadings= average±2*S.E.M.)

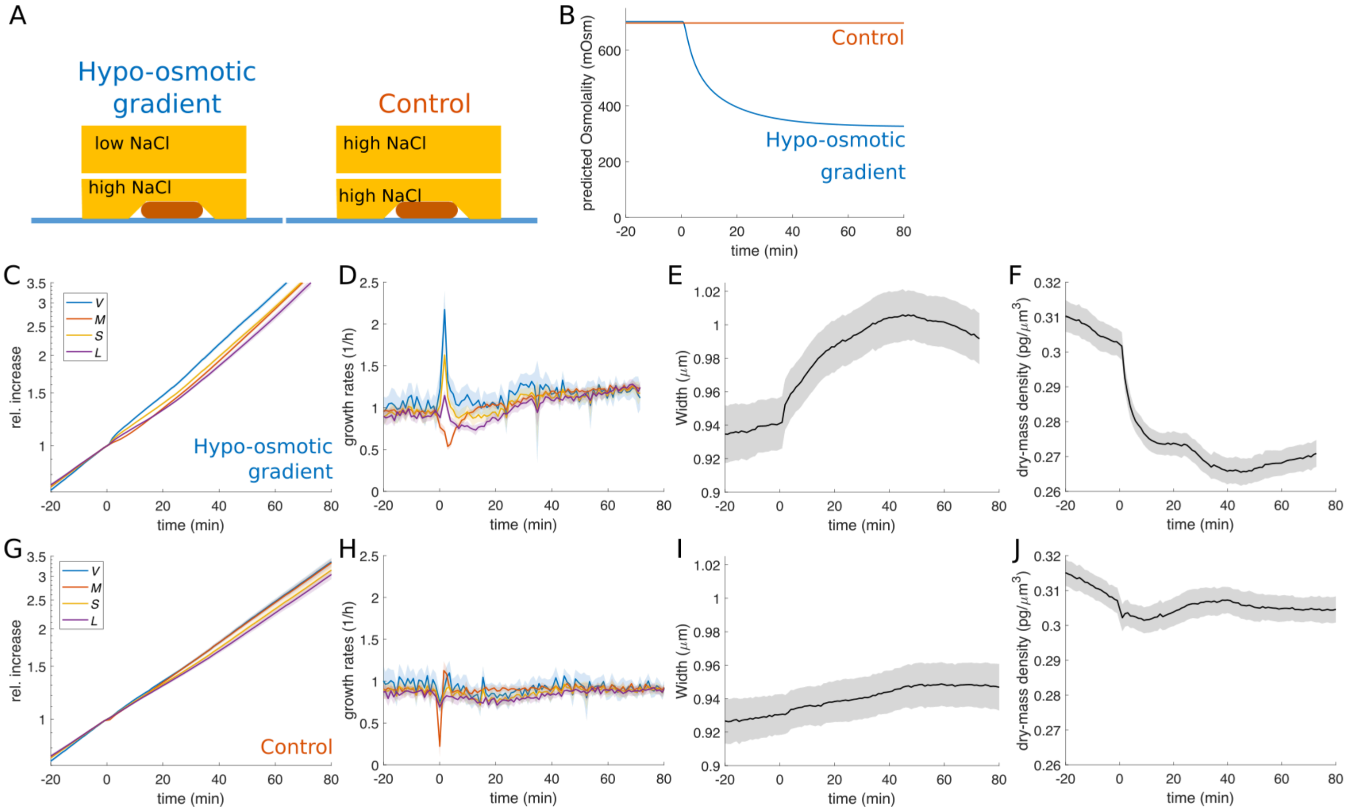

**Fig. S14. Hypoosmotic ramp leads to similar changes of cell dimensions as nutrient upshifts.**
**A-B:** Cartoon of hypo-osmotic gradient experiment and control. An agar pad with low NaCl is placed on top of a microscopy pad with high NaCl, leading to a time-dependent reduction of osmolality (B), calculated from a 1D diffusion model. **C-J:** Hypo-osmotic gradient experiment (C-F), and control experiment (G-J). Relative change of volume, mass, surface area, length (C,G), relative rates (D,H), width (E,I), and mass density (F,J) as a function of time.

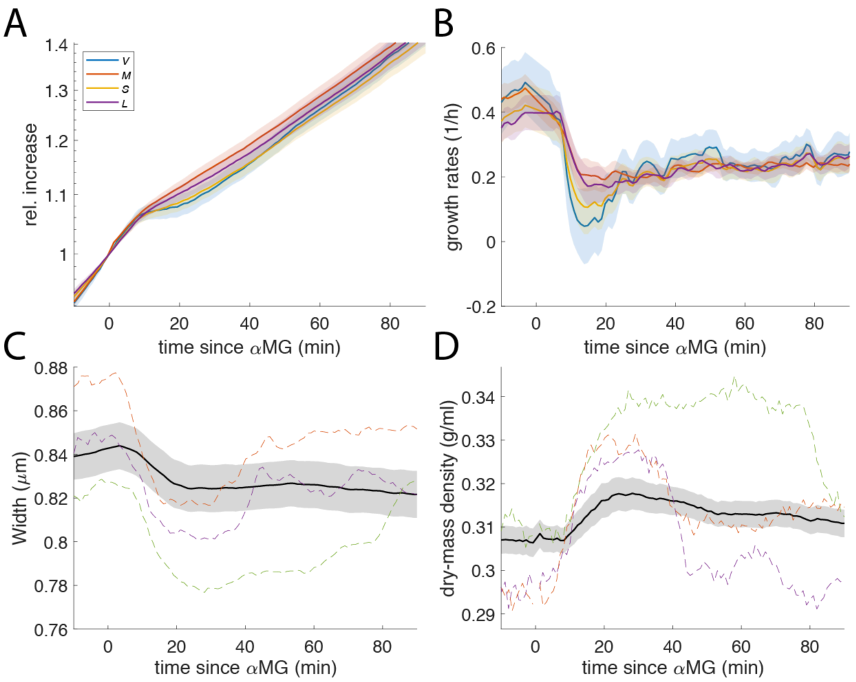

**Fig. S15. Depletion of glucose using αMG.** Relative change of volume, mass, surface area, length (**A**), relative rates (**B**), width (**C**), and mass density (**D**) as a function of time during application of alpha-MG. (solid lines+shadings= average±2*S.E.M.; colored dashed lines=single-cell traces)

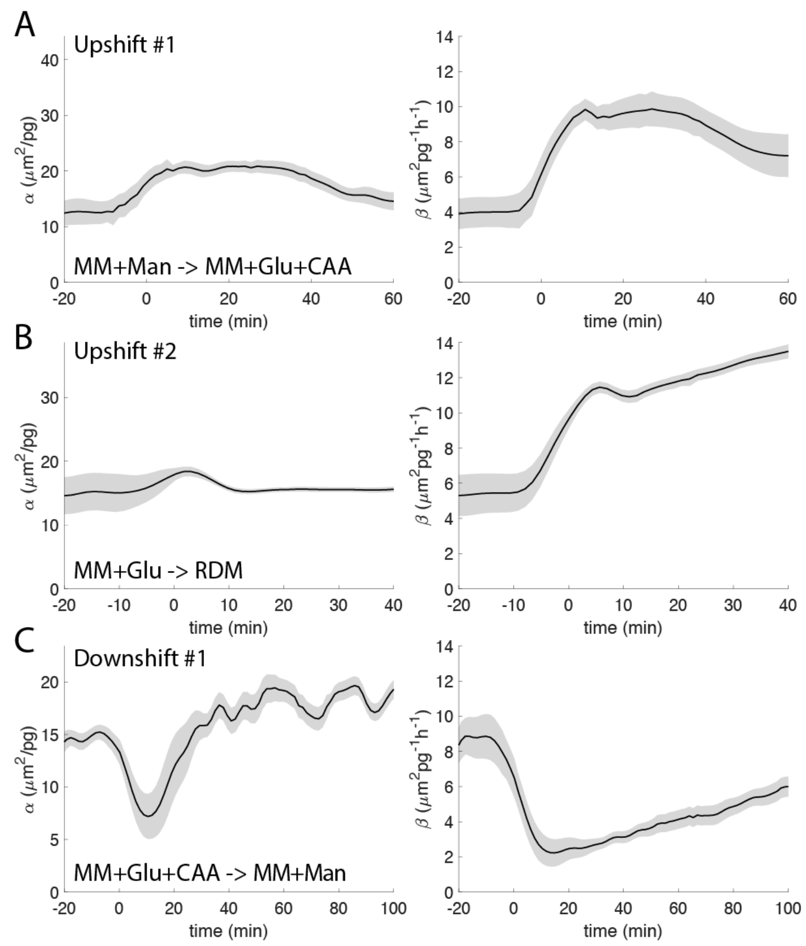

**Fig. S16. Comparing different surface-to-mass coupling constants during nutrient shifts.** Time-dependency of the coupling constants $\alpha = \mathrm{d}S/\mathrm{dM}$ (left panels) and $\beta = 1/M(\mathrm{d}S/\mathrm{d}t)$ (right panels) during the three nutrient shifts applied to filamenting cells (S290) in flow chambers presented in Fig. 3B,C,E. Apart from transient changes due to Turgor right after the shift, $\alpha$ varies little before and after the shift, while $\beta$ changes in a growth-rate dependent manner, nearly proportionally to mass growth rate.

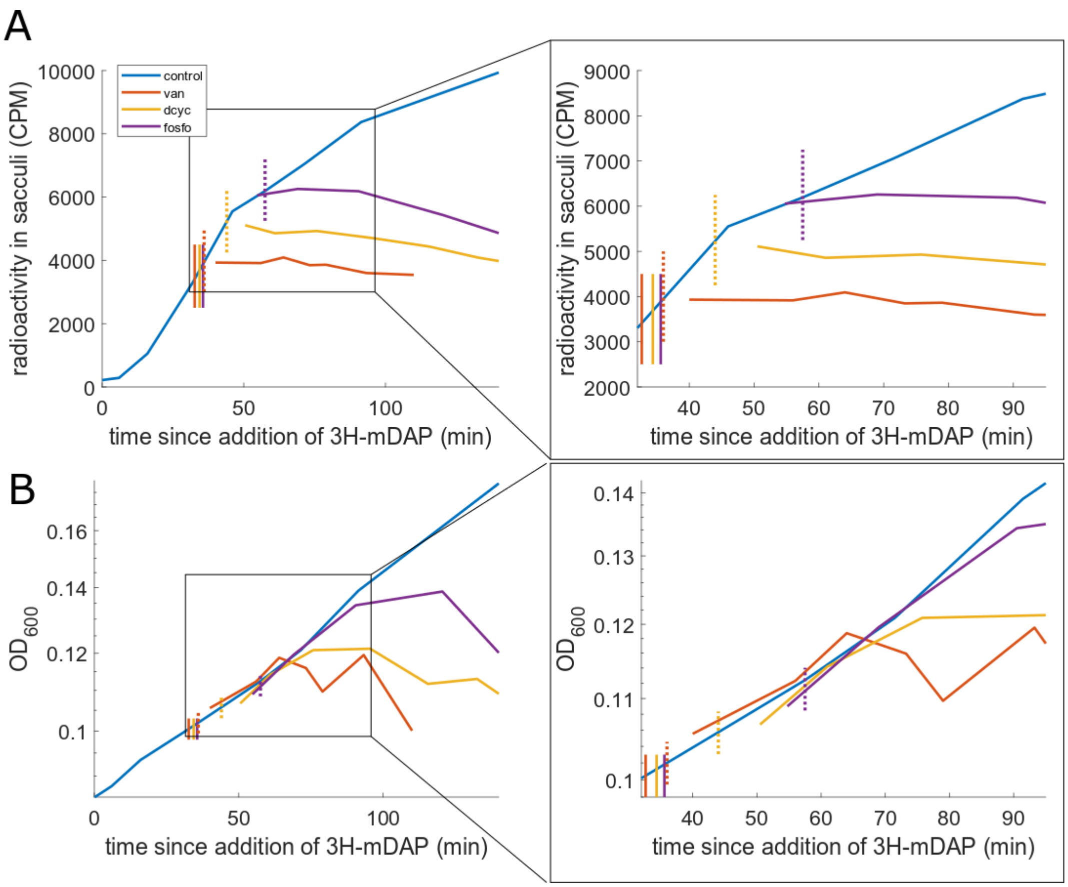

**Fig. S17. Measurements of cell-wall synthesis with 3H-mDAP labeling.** Increase of radioactivity in isolated cell-wall sacculi after addition of 3H-mDAP (**A**) and $OD_{600}$ measurements (**B**) for an untreated control and for treatments of vancomycin (100ug/ml), D-cycloserine (1mM), and Fosfomycin (500ug/ml). Solid vertical lines indicate the time of drug addition, dotted vertical lines indicate the time of precursor-incorporation stop (estimated as the time when the control reaches the maximum signal of the drug-treated samples), boxes indicate zoom-in shown on the right. 3H-mDAP incorporation stops after ~3min (van), ~10min (dcyc), and ~22min (fosfo). After mDAP-incorporation arrest, $OD_{600}$ increased in an unperturbed fashion by about 10-20%, before growth curves of drug-treated cultures started to deviate from the control.

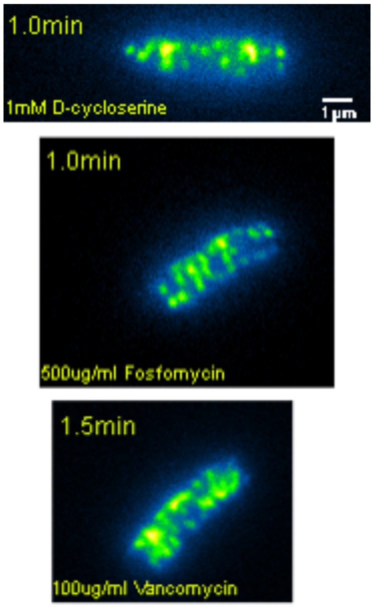

**Movie S1-3. MreB-msfGFP motion arrest upon cell-wall drug treatments.** Movies of MreB-msfGFP in cells growing on agar pads during treatment with D-cycloserine (S1), Fosfomycin (S2), or Vancomycin (S3). Each slice of the multi-dimensional TIFF stack contains a 20 s-long movie of the same cell taken at different time points after drug addition (time points 7-10 min apart, as indicated in the movies). Strains used were S257 (D-cycloserine, Fosfomycin), S382 (Vancomycin).
