## Supplementary figures and images for "Bacteria control cell volume by coupling cell-surface expansion to dry-mass growth"

### MovieS1.tif

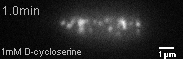

### MovieS2.tif

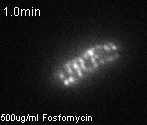

### MovieS3.tif

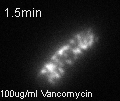
